# Scientifically Uninformed Legislation System Undermines Ecological Sustainability of Yangtze River

**DOI:** 10.1101/2025.07.11.663308

**Authors:** Binbin Wang, Hao Zhuang, Jason H. Knouft

## Abstract

Legal frameworks typically aim to protect environmental systems through regulations, treaties, and conservation measures. However, without scientifically informed enforcement, these policies may inadvertently exacerbate environmental degradation, particularly when addressing complex socio-environmental issues. We reviewed the development of China’s environmental legislation related to ecosystem protection in the Yangtze River watershed and evaluated its effectiveness by analyzing illegal fishing cases from 2015 to 2021, prosecuted under China’s Environmental Public Interest Litigation framework. Our findings reveal that court-ordered fish release remedies often undermine, rather than support, the sustainability of the Yangtze River ecosystem. This is primarily due to mismatches between the taxonomic identity of illegally caught fishes and those specified for release, species misidentification, and a bias toward economically valuable species in court-ordered release decisions. These unintended discrepancies between environmental protection goals and actual ecological outcomes underscore the need to integrate scientific expertise into legal practice to enhance the sustainability of socio-environmental systems.

## Introduction

The sustainability of freshwater systems is fundamentally important to society and interwoven into all aspects of the United Nations 17 Sustainable Development Goals. However, increased pressures on freshwater fisheries have compromised the sustainability of freshwater ecosystems globally (FAO, 2016). Legal frameworks have been implemented to safeguard these systems through regulations, treaties, and conservation measures (Bell et al. 2013, Yang & Percival 2009). Nevertheless, the complexity of freshwater systems and their interconnectedness with human systems pose significant challenges to the effective implementation and enforcement of these legal frameworks. Factors such as limited resources, conflicting interests, and a lack of scientifically grounded remediation strategies can inadvertently undermine the sustainability of these socio-environmental systems.

The Yangtze River basin, one of the world’s largest freshwater ecosystems, supports the highest freshwater fish diversity in China (Chen et al. 2009) and contributes up to 60% of the national wild freshwater fisheries catch (Yu 1988). As such, the basin plays a crucial role in China’s food security, economy, and employment. However, wild fish production in the Yangtze River basin has steadily declined since peaking in the 1950s, reaching just 27.3% of its former levels by 2021, which is concurrant with significant loss of biodiversity (Zhang et al. 2020). Overfishing is one of the major drivers of this decline (Ye et al. 2014, Huang and Li 2016), despite efforts by the Chinese government over the past two decades to address the issue through various policies targeting overfishing in the basin (Chen et al., 2020).

A seasonal fishing moratorium (April – June) was implemented in 2002 on the Yangtze River and escalated to a “10-year moratorium on commercial fishing” by the Ministry of Agriculture and Rural Affairs in 2021 (http://www.cjyzbgs.moa.gov.cn/tzgg/201912/t20191227_6334009.htm). The ban prohibits all commercial fishing in both the mainstem and tributaries of the watershed, with local government agencies tasked with strengthening law enforcement capacity. Furthermore, the Yangtze River Protection Law (YRPL, https://english.mee.gov.cn/Resources/laws/environmental_laws/202104/t20210407_827604.shtm l), the first-ever river-watershed protection legislation in China, was approved by the National People’s Congress in December 2020 and came into effect in March 2021. The YRPL calls for the establishment of a National Yangtze River Watershed Coordination Mechanism and places obligations on national agencies and provincial governments to align with the goals of ecological protection and improved water quality. Nevertheless, illegal, unreported, and unregulated (IUU) fishing appears to have increased across the watershed since the implementation of the seasonal fishing moratorium in 2002, and the problem has become even more pronounced following the implementation of the 10-year ban on commercial fishing throughout the watershed in 2021 (Zhang et al 2020; Jin et al. 2022).

The conventional “command-and-control” approach dominated by public law enforcers in China is not sufficient or effective in controlling IUU fishing (Mei et al. 2018), due to local shortages of staff and limited capacity within public agencies. Therefore, in 2015, a new legal policy, the Environment Public Interest Litigation (EPIL), was enacted in China, which grants standing right to registered civil society organizations to sue individuals, private entities, and government agencies for environmental misconduct to protect the “public interest” (Zhuang and Wolf, 2021). The policy is supported by two major legislative provisions: the Amendment of The Civil Procedure Law of China (Article 55 2012) and the Revision of the Environmental Protection Law of China (Article 58 2014), which define the eligibility and rights for civil society organizations in practicing law. This policy marks the first time the Chinese judicial system has incorporated a civil component into environmental law enforcement, alongside public prosecutors, police, and courts. This addition has encouraged public prosecutors to take a more active role in enforcing environmental laws (Zhuang & Wolf 2021).

During the first seven years of the EPIL, there was a steady increase in legal cases focusing on IUU fishing, along with a wide range of environmental issues including water, soil and air pollution, deforestation, illegal mining, biodiversity loss, and climate change. While the IUU fishing cases have applied novel judicial remedies to restore fish species and support ecosystem sustainability in the Yangtze River basin, the associated litigation procedures and remediation judgments reveal a disconnect between legal practices and scientifically informed management.

In this study, we compiled empirical legal data from EPIL cases over the first seven years of the program (2015-2021) across three major provinces within the Yangtze River basin. We then evaluated whether court-imposed remedies aimed at mitigating IUU fishing were appropriate in relation to the goals of conserving biodiversity and maintaining ecosystem integrity. Based on these assessments, we discuss the critical role of science in informing legal practice, an integration that can ultimately support the development of sustainable social-environmental systems.

## Results

### Implementation of the EPIL Policy in China

EPIL cases increased dramatically across China from 2015 - 2021 (Fig. 1a), with the most recent official report issued by the Supreme Court of China indicating that 23,000 cases were filed from January 2018 to September 2023. We reviewed the EPIL cases ruled by local courts in three provinces (Hubei, Sichuan, and Jiangsu) along the Yangtze River and found 516 cases from 2015 to 2021 covering various environmental themes (Fig. 2). Among these cases, 120 involved illegal fishing (all filed by the Prosecutors). The number of illegal fishing cases increased from 2015 through 2019, then declined in 2020 and 2021 (Fig. 1b), possibly due to the reduced fishing activity or diminished legal oversight during the COVID-19 pandemic (Knouft 2022). The number of cases was higher in Hubei and Sichuan, and lower in Jiangsu. Additionally, 68 illegal fishing cases occurred in the mainstem of the Yangtze River, 39 cases in the tributaries, and eight cases in the lakes adjacent to the river system, while five cases did not identify specific locations (Fig. 1c). Most cases in Sichuan Province (78%) were in the tributaries, while most cases in Hubei Provice (82%) and Jiangsu Province (67%) were in the mainstem of the Yangtze River.

**Figure 1.**
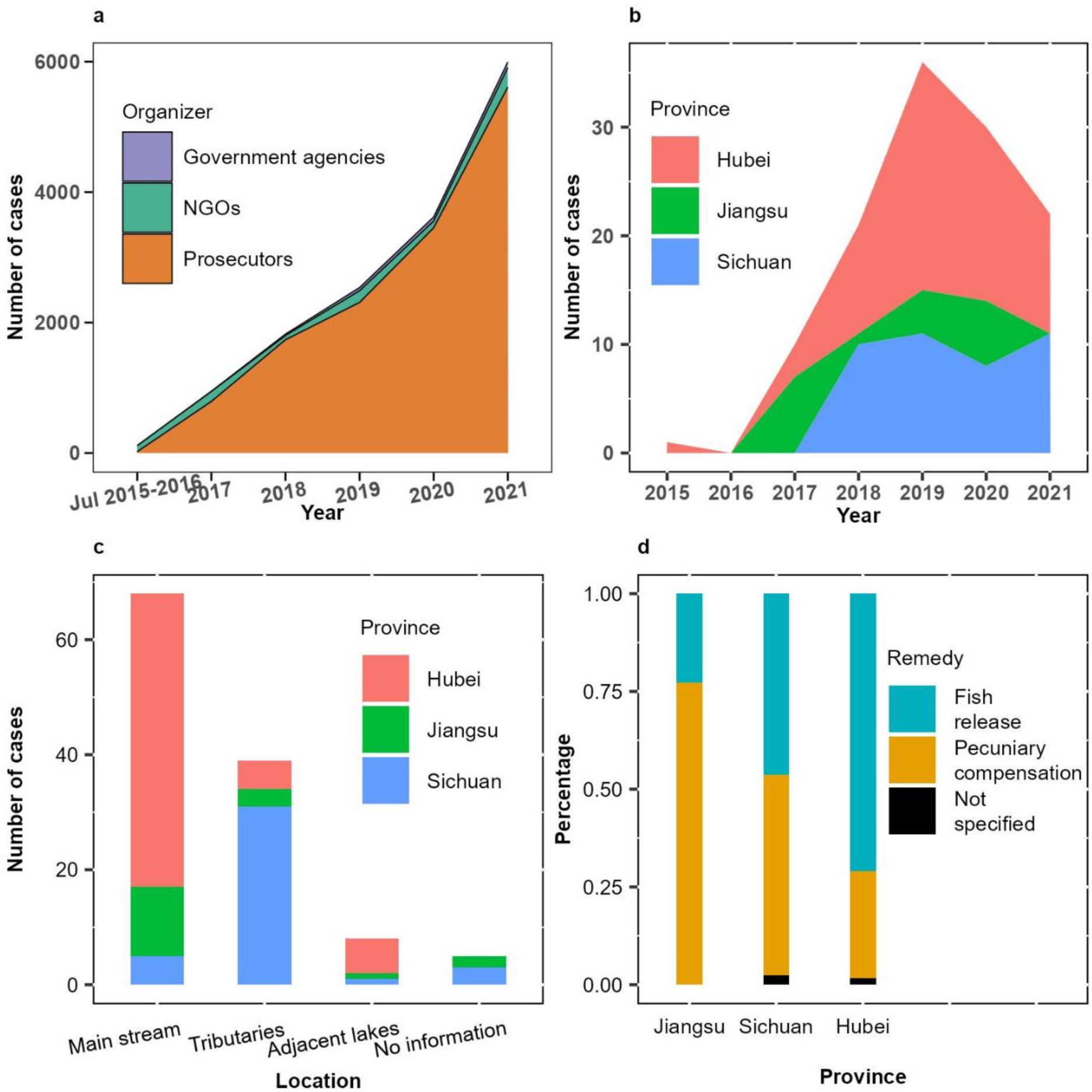
Summary of Environmental Public Interest Litigation (EPIL) and illegal fishing cases in the three study provinces along the Yangtze River. a. Temporal trends in total EPIL cases from 2015 to 2021, categorized by enforcement agencies. b. Number of illegal fishing EPIL cases by province. c. Spatial distribution of illegal fishing case locations across Sichuan, Hubei, and Jiangsu Provinces. d. Proportional use of different environmental restoration remedies in illegal fishing cases.

**Figure 2.**
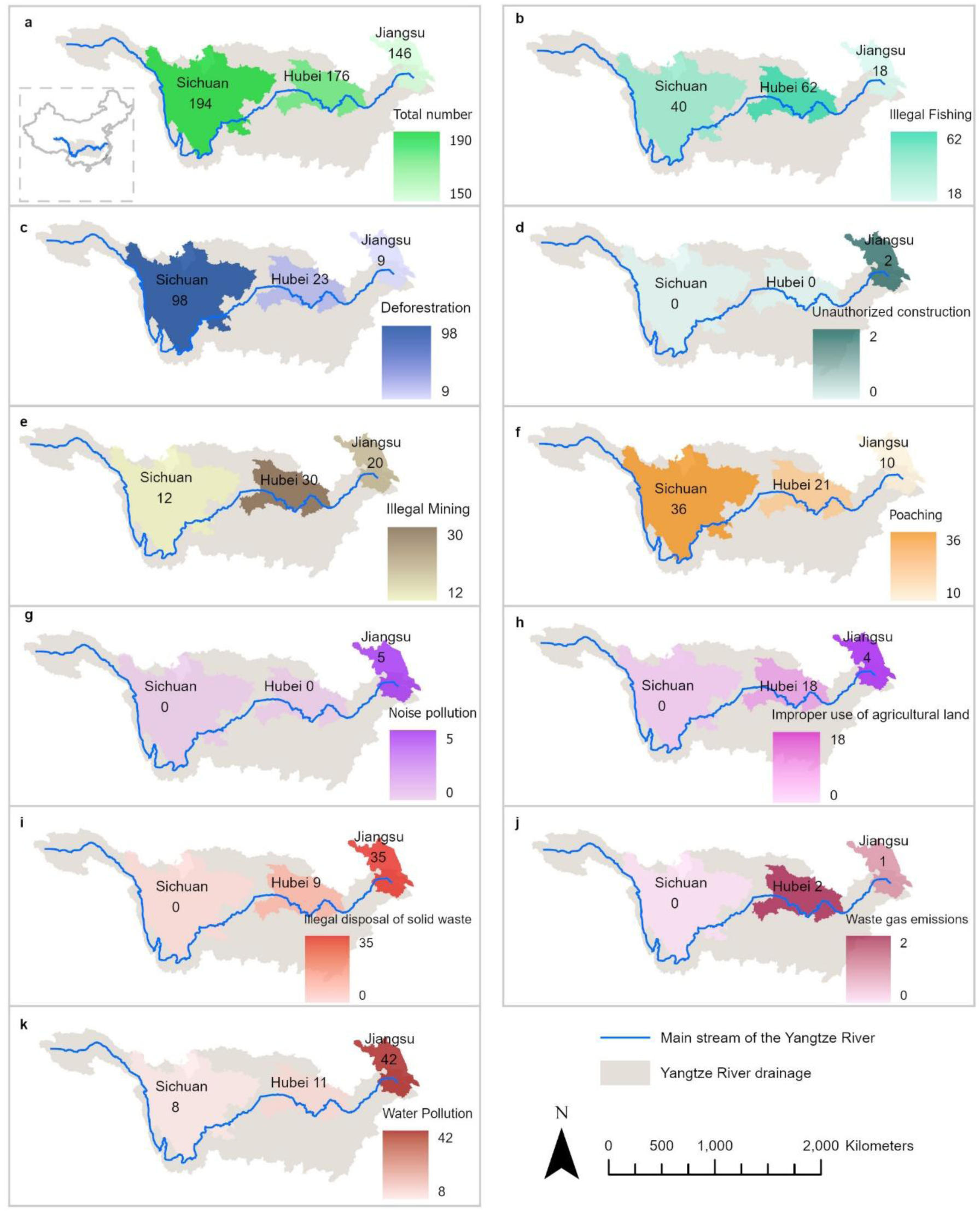
Distribution of EPIL case types across the three study provinces along the Yangtze River. The numbers in the graphs are total numbers and categorical breakdowns of EPIL cases from 2015 to 2021 in Sichuan, Hubei, and Jiangsu Provinces.

### Fish release as an approach to remediation

In each case, local courts recommended either fish release or pecuniary compensation (Fig. 1d). Fish released were purchased from hatcheries as ordered by the court, while pecuniary compensation was determined based on the economic value of the species and the potential damage to local fisheries. Hubei and Sichuan provinces reported higher frequencies of fish release compared to Jiangsu Province.

A total of 46 fish taxa were identified in all cases that reported illegal captures, with 43 taxa identified to species-level and three to genus-level or subfamily-level (Table 1). In total, 9,483 kg and 117,675 individuals were caught in cases with known catch quantities, with 4,123 kg and 117,670 individuals identified to some taxonomic level (Table S1). As required by court ruling, 26 fish taxa were released, with 24 identified to the species-level, one to the genus-level, and one to the subfamily-level (Table S2). In total, 12,394 kg of adult fishes plus 5,706,324 individual juvenile fishes were released, including 2,120 kg of adults and 2,390,499 juveniles identified to some taxonomic level. Only 21 of the 43 species caught were released as court-ordered. Excluding the non-native *Oreochromis sp.*, 24 native taxa (22 species + 2 genera) were caught and not released during these reintroduction efforts. Additionally, there were three species released but not caught during illegal fishing (Table 1).

**Table 1.**
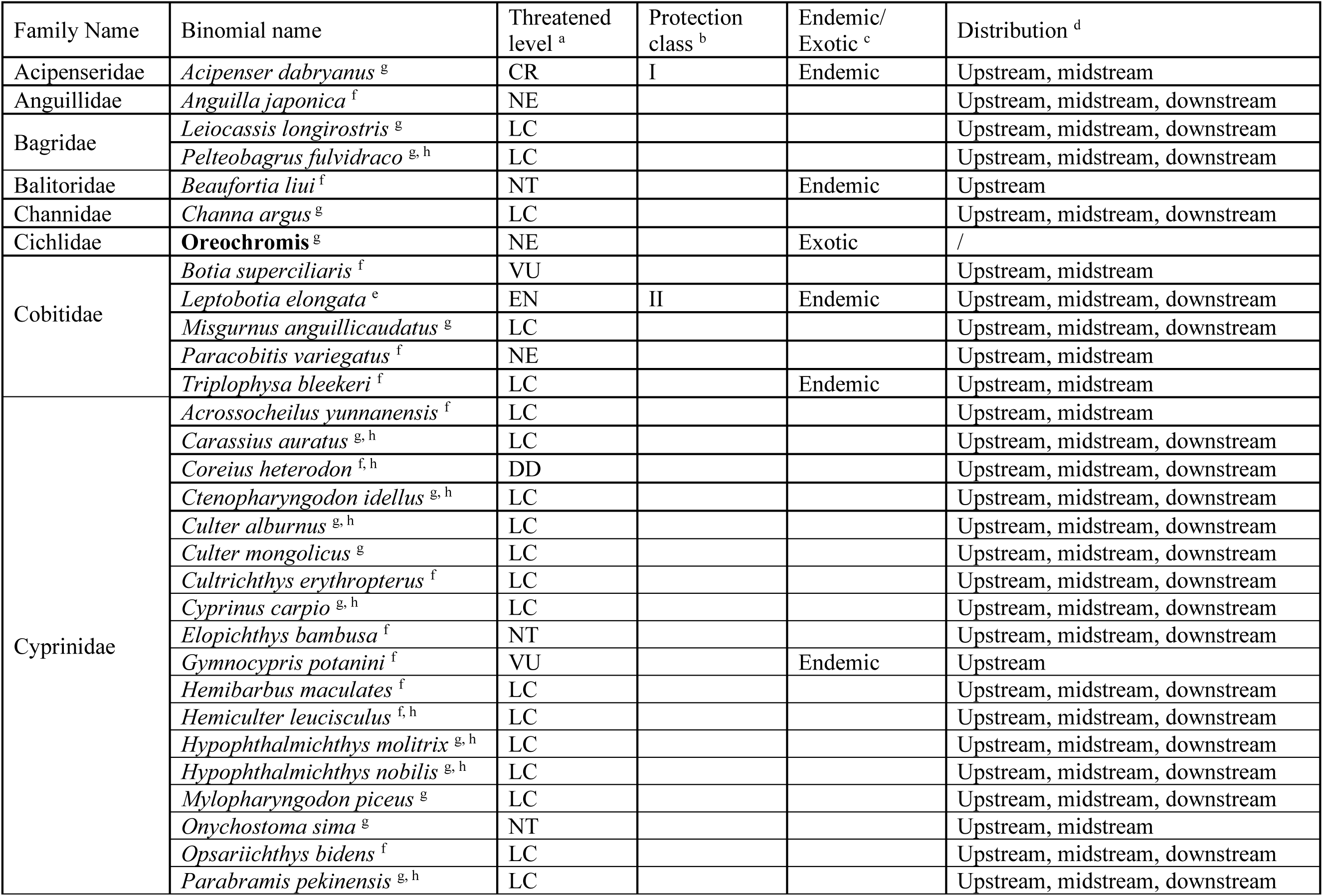
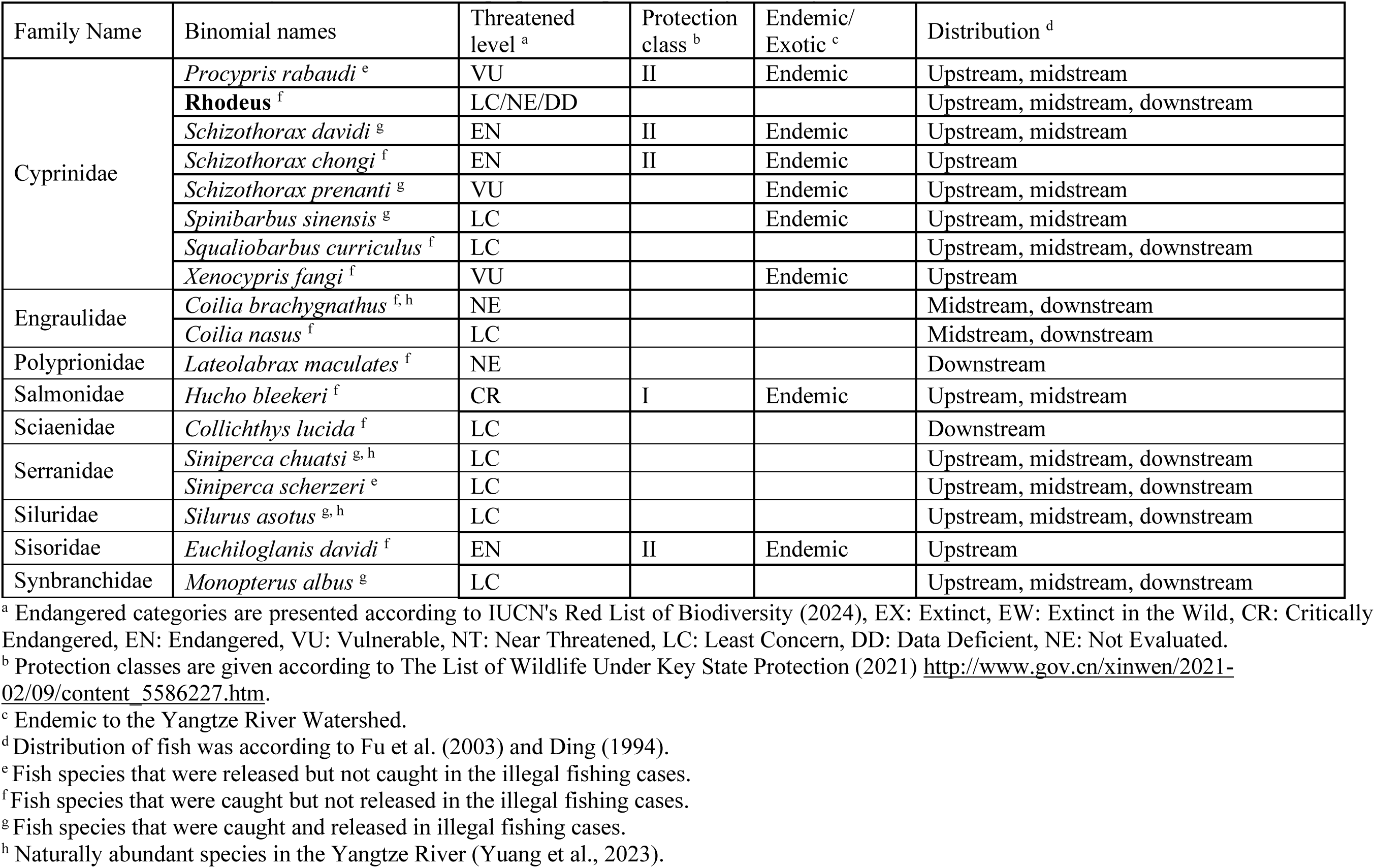
Summary of fish taxa and their properties reported in illegal fishing cases.

| Family Name | Binomial name | Threatened level <sup>a</sup> | Protection class <sup>b</sup> | Endemic/ Exotic <sup>c</sup> | Distribution <sup>d</sup> |
| --- | --- | --- | --- | --- | --- |
| Acipenseridae | *Acipenser dabryanus* <sup>g</sup> | CR | I | Endemic | Upstream, midstream |
| Anguillidae | *Anguilla japonica* <sup>f</sup> | NE |  |  | Upstream, midstream, downstream |
| Bagridae | *Leiocassis longirostris* <sup>g</sup> | LC |  |  | Upstream, midstream, downstream |
|  | *Pelteobagrus fulvidraco* <sup>g, h</sup> | LC |  |  | Upstream, midstream, downstream |
| Balitoridae | *Beaufortia liui* <sup>f</sup> | NT |  | Endemic | Upstream |
| Channidae | *Channa argus* <sup>g</sup> | LC |  |  | Upstream, midstream, downstream |
| Cichlidae | **Oreochromis** <sup>g</sup> | NE |  | Exotic | / |
| Cobitidae | *Botia superciliaris* <sup>f</sup> | VU |  |  | Upstream, midstream |
|  | *Leptobotia elongata* <sup>e</sup> | EN | II | Endemic | Upstream, midstream, downstream |
|  | *Misgurnus anguillicaudatus* <sup>g</sup> | LC |  |  | Upstream, midstream, downstream |
|  | *Paracobitis variegatus* <sup>f</sup> | NE |  |  | Upstream, midstream |
|  | *Triplophysa bleekeri* <sup>f</sup> | LC |  | Endemic | Upstream, midstream |
| Cyprinidae | *Acrossocheilus yunnanensis* <sup>f</sup> | LC |  |  | Upstream, midstream |
|  | *Carassius auratus* <sup>g, h</sup> | LC |  |  | Upstream, midstream, downstream |
|  | *Coreius heterodon* <sup>f, h</sup> | DD |  |  | Upstream, midstream, downstream |
|  | *Ctenopharyngodon idellus* <sup>g, h</sup> | LC |  |  | Upstream, midstream, downstream |
|  | *Culter alburnus* <sup>g, h</sup> | LC |  |  | Upstream, midstream, downstream |
|  | *Culter mongolicus* <sup>g</sup> | LC |  |  | Upstream, midstream, downstream |
|  | *Cultrichthys erythropterus* <sup>f</sup> | LC |  |  | Upstream, midstream, downstream |
|  | *Cyprinus carpio* <sup>g, h</sup> | LC |  |  | Upstream, midstream, downstream |
|  | *Elopichthys bambusa* <sup>f</sup> | NT |  |  | Upstream, midstream, downstream |
|  | *Gymnocypris potanini* <sup>f</sup> | VU |  | Endemic | Upstream |
|  | *Hemibarbus maculatus* <sup>f</sup> | LC |  |  | Upstream, midstream, downstream |
|  | *Hemiculter leucisculus* <sup>f, h</sup> | LC |  |  | Upstream, midstream, downstream |
|  | *Hypophthalmichthys molitrix* <sup>g, h</sup> | LC |  |  | Upstream, midstream, downstream |
|  | *Hypophthalmichthys nobilis* <sup>g, h</sup> | LC |  |  | Upstream, midstream, downstream |
|  | *Mylopharyngodon piceus* <sup>g</sup> | LC |  |  | Upstream, midstream, downstream |
|  | *Onychostoma sima* <sup>g</sup> | NT |  |  | Upstream, midstream |
|  | *Opsariichthys bidens* <sup>f</sup> | LC |  |  | Upstream, midstream, downstream |
|  | *Parabramis pekinensis* <sup>g, h</sup> | LC |  |  | Upstream, midstream, downstream |

Table 1 (continued) Summary of fish taxa and their properties reported in illegal fishing cases.
| Family Name | Binomial names | Threatened level <sup>a</sup> | Protection class <sup>b</sup> | Endemic/ Exotic <sup>c</sup> | Distribution <sup>d</sup> |
| --- | --- | --- | --- | --- | --- |
| Cyprinidae | *Procypris rabaudi* <sup>e</sup> | VU | II | Endemic | Upstream, midstream |
|  | **Rhodeus** <sup>f</sup> | LC/NE/DD |  |  | Upstream, midstream, downstream |
|  | *Schizothorax davidi* <sup>g</sup> | EN | II | Endemic | Upstream, midstream |
|  | *Schizothorax chongi* <sup>f</sup> | EN | II | Endemic | Upstream |
|  | *Schizothorax prenanti* <sup>g</sup> | VU |  | Endemic | Upstream, midstream |
|  | *Spinibarbus sinensis* <sup>g</sup> | LC |  | Endemic | Upstream, midstream |
|  | *Squaliobarbus curriculus* <sup>f</sup> | LC |  |  | Upstream, midstream, downstream |
|  | *Xenocypris fangi* <sup>f</sup> | VU |  | Endemic | Upstream |
| Engraulidae | *Coilia brachygnathus* <sup>f, h</sup> | NE |  |  | Midstream, downstream |
|  | *Coilia nasus* <sup>f</sup> | LC |  |  | Midstream, downstream |
| Polyprionidae | *Lateolabrax maculatus* <sup>f</sup> | NE |  |  | Downstream |
| Salmonidae | *Hucho bleekeri* <sup>f</sup> | CR | I | Endemic | Upstream, midstream |
| Sciaenidae | *Collichthys lucida* <sup>f</sup> | LC |  |  | Downstream |
| Serranidae | *Siniperca chuatsi* <sup>g, h</sup> | LC |  |  | Upstream, midstream, downstream |
|  | *Siniperca scherzeri* <sup>e</sup> | LC |  |  | Upstream, midstream, downstream |
| Siluridae | *Silurus asotus* <sup>g, h</sup> | LC |  |  | Upstream, midstream, downstream |
| Sisoridae | *Euchiloglanis davidi* <sup>f</sup> | EN | II | Endemic | Upstream |
| Synbranchidae | *Monopterus albus* <sup>g</sup> | LC |  |  | Upstream, midstream, downstream |

<sup>a</sup> Endangered categories are presented according to IUCN's Red List of Biodiversity (2024), EX: Extinct, EW: Extinct in the Wild, CR: Critically Endangered, EN: Endangered, VU: Vulnerable, NT: Near Threatened, LC: Least Concern, DD: Data Deficient, NE: Not Evaluated.
<sup>b</sup> Protection classes are given according to The List of Wildlife Under Key State Protection (2021) http://www.gov.cn/xinwen/2021-02/09/content_5586227.htm.
<sup>c</sup> Endemic to the Yangtze River Watershed.
<sup>d</sup> Distribution of fish was according to Fu et al. (2003) and Ding (1994).
<sup>e</sup> Fish species that were released but not caught in the illegal fishing cases.
<sup>f</sup> Fish species that were caught but not released in the illegal fishing cases.
<sup>g</sup> Fish species that were caught and released in illegal fishing cases.
<sup>h</sup> Naturally abundant species in the Yangtze River (Yuang et al., 2023).

#### 1) Choice of the fish species for release

To better demonstrate the impact of the law enforcement practices on the fish communities in the Yangtze River basin, we grouped the fish species identified in the EPIL cases into three categories: Common Aquaculture Species (CAS), Vulnerable and Endangered Species (VES), and Naturally Abundant Species (NAS) (definitions provided in the Methods section).

The relative frequency of CAS in the Yangtze River system were likely increased due to court-ordered releases, as these species were more frequently released than illegally caught. The 14 CAS accounted for 30.4% of the 46 illegally caught taxa and 63.9% of all capture instances (i.e., each time a taxon was caught in a case, it was counted as one capture instance). However, CAS made up 58.3% of the species released under court orders and 87.2% of all release instances (i.e., each time a taxon was released in a case, it was counted as one release instance). Notably, the annual aquaculture production of the 14 CAS in these three provinces was more strongly correlated with their release instance than with their caught instance (p=0.009, Extened Fig. 1). On the contrary, the status of VES may have been weakened due to their relatively lower likelihood of being released compared to being caught. Ten VES accounted for 6.9% of capture instances, but seven of them were not released, representing 17.9% instances where a species was caught but not released. In total, nine threatened species and five nationally protected species were caught, yet 67% and 60% of these species, respectively, were not included in the court-order releases. The status of non-CAS and non-VES species was also weakened. Specifically, 17 (68%) of the 25 species caught but not released were non-CAS and non-VES, while only five (28.8%) of the 24 released species were non-CAS and non-VES species, accounting for 9.4% of release instances. Additionally, the five released non-CAS and non-VES (*Chanodichthys mongolicus*, *Onychostoma sima*, *Siniperca scherzeri*, *Culter alburnus*, and *Spinibarbus sinensis*) are frequently reported as important economic species to be released into the Yangtze River basin, even without occureance of illegal fishing or court orders (Deng et al. 2010, Li et al 2013, Wang et al. 2015, Jiang et al. 2017, Xie et al. 2019, Ren et al. 2020).

#### 2) Substitution of fish taxa in court-ordered releases

The goal of court-ordered fish releases is to remdiate the impacts of illegal fishing; however, mismatches occurred between the fish taxa caught and those released in 18 of the 21 cases with known taxa. In 13 of these cases, the caught taxa were substituted with new taxa during release; two cases released more taxa than were caught; three cases released fewer taxa than were caught; and only three cases released the same taxa as were caught (Table S3).

Twenty-two species were involved in the mismatched cases (Fig. 3, Table S3), including 11 CAS and 4 VES. In the mismatched cases, CAS were highly likely (81.5%) to be released as substitutes for other species or released without being initially caught, while the VES were more likely (60%) to be substituted by other species or caught without being released. Specifically, *Hypophthalmichthys molitrix* was released most frequently (38.9% of cases), followed by *Ctenopharyngodon idella* and *Cyprinus rubrofuscus* (22.2% of cases each) (Fig. 3). These three species were among the top five in aquaculture production and accounted for 45.8% of the total freshwater aquaculture production in these three provinces (Table S5). In summary, CAS were inappropriately oversupplemented, while VES and other species were inappropriately underrepresented in the mismatched releases (Fig. 3).

**Figure 3.**
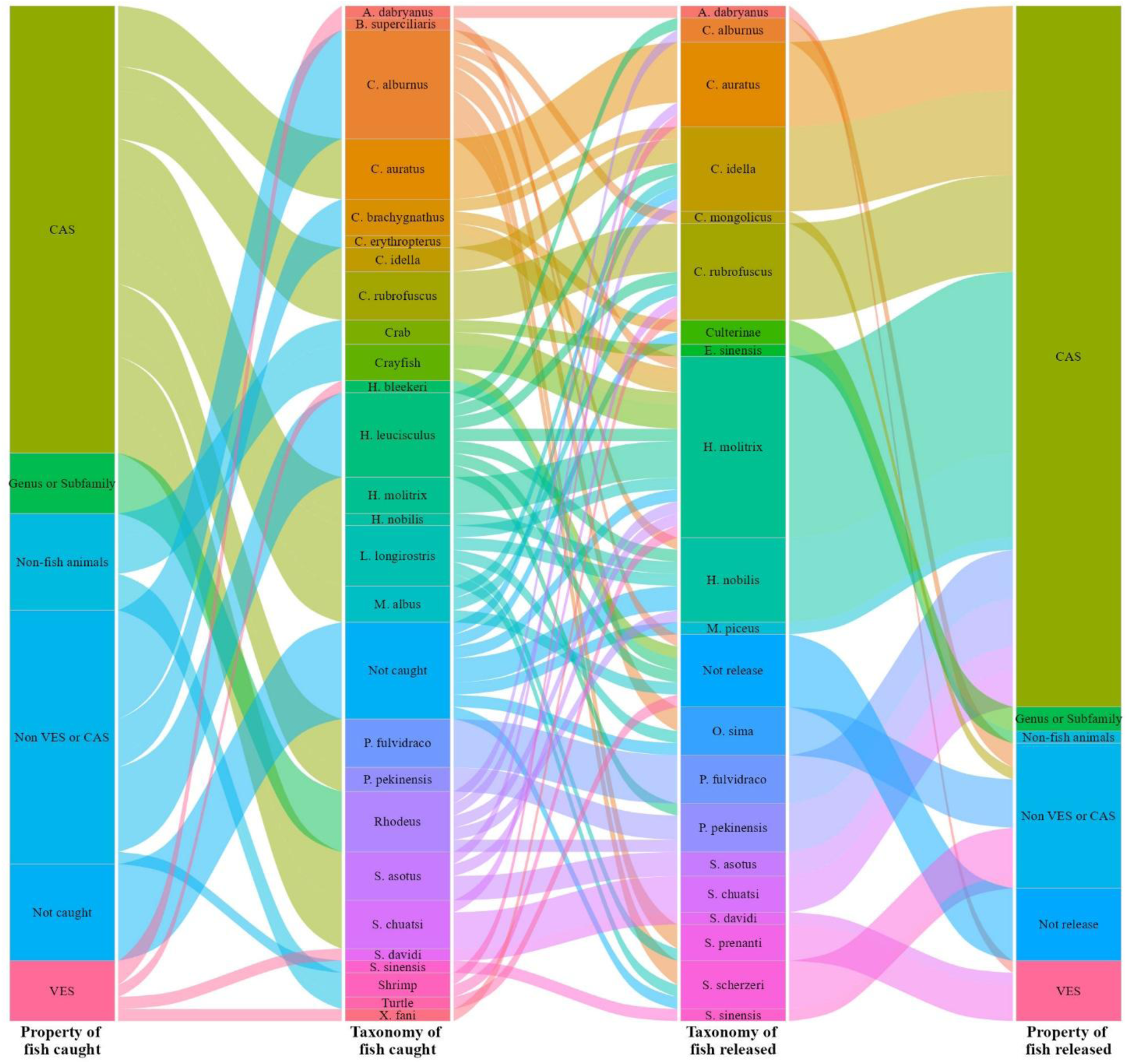
Taxonomic replacement patterns in illegal fishing cases (n=21) with identified taxonomic names for both caught and released fish. Strata heights in the first two columns represent relative release instances attributed to each taxonomic category and property of the originally caught taxa. Strata in the third and fourth columns represent relative release instances by taxonomic category and properity of the substitute (released) taxa. The width of each flow between caught and released taxa is proportional to the number of instances in which a caught taxon was replaced with the same or a different taxon in the same case.

#### 3) Inaccurate identification of fish species

In some cases, inaccurate identification of illegally captured fish species and a lack of recognition of the different ecological roles of similar species occurred. For example, 295.5 kg of *Coilia brachygnathus* (Yangtze Grenadier Anchovy), caught in Long Lake, a lake adjacent to the middle Yangtze River, were misidentified as *Coilia mystus,* an anadromous species that rarely migrates to the middle reaches of the Yangtze River (Xu et al. 2014). In contrast, *Coilia brachygnathus* is a dominant species and relatively abundant in the middle Yangtze River (Yang et al. 2023). Additionally, the use of common names instead of scientific names resulted in inaccurate identifications, as multiple common names are used for the same species in different regions along the Yangtze River (Ding 1994). For example, *Hemiculter leucisculus* was referred to using six different common names in the reviewed cases.

## Discussion

The increases in the number of EPIL cases from 2015 through 2021 reflects the rapid growth of Non-Governmental Organization (NGO) and Public Prosecutors’ enforcement efforts in public-interest litigation (Fig. 1a). In early 2017, public prosecutors were granted exclusive standing to file litigation against government agencies (administrative litigation) (Civil Procedural Law Article 55, Administrative Procedural Law Article 25). Since then, the public prosecutor has become the driving force behind EPIL implementation. Environmental restoration is the primary goal among the judicial remedies commonly implemented in EPIL cases, which aligns with China’s national "environmental protection" strategy, emphasizing increased investment in environmental improvement and restoration (Master Plan of Environmental Protection and Environmental Restoration 2021-2035, https://www.ndrc.gov.cn/xxgk/zcfb/tz/202006/P020200611354032680531.pdf). Fish release and tree planting are frequently used as environmental restoration approaches in EPIL court judgements, particularly in prosecutors’ cases. However, the judicial processes in these cases often lack systematic guidance for restoration practices, particularly fish releases, rendering remediation efforts inappropriate and harmful to natural systems.

### Lack of science-based consultation in the judicial system practices

EPIL has a clear goal of protecting ecological integrity and enhancing environmental quality. To achieve this goal, law enforcement must be based on an accuate understanding of environmental science and ecological conservation. This also requires supporting law enforcement (i.e., police, prosecutors, NGO practitioners and court judges at decentralized levels) with accurate biological and environmental knowledge. However, local police officers are not trained in environmental issues, as their primary mandates are focused on investigating criminal actions. Local court judges and public prosecutors have been involved in the EPIL system since 2015 and 2017, respectively, but the conventional mandates and knowledge structure of the judiciary staff do not include expertise in environmental science, such as fish identification. As a result, there is a gap in staff capacity, expertise, and knowledge, which can lead to practical errors, such as misidentification of fish taxa, mismatches between caught and released fish species, and bias toward economically significant species, as discussed above.

In some cases, however, local courts have effectively relied on expert advice to assess the extent of environmental damage caused by misconduct, recommend appropriate restoration methods, such as fish releases, and monitor the implementation of court judgments. The selection of experts or institutions with experience in fisheries science varies widely among the courts in our database. Experts may come from leading research institutes specializing in fisheries, such as the Yangtze River Fisheries Research Institute and the Freshwater Fisheries Research Center of the Chinese Academy of Fisheries Sciences. Experts may also be legal professionals from government-affiliated organizations, such as the Justice Authenticate Center or Forest Forensic Appraisal Centers. Additionally, some local courts consult the local agriculture bureau or agricultural extension stations, given that fisheries are often managed within China’s agricultural sector. It is also common to create an expert pool that includes well-recognized researchers and specialists in the field. For example, in a case from Hubei, a fish breeding center was invited to serve as an appraisal party in litigation.

While efforts have been made to connect informed scientists with court-ordered releases, these efforts have not provided the broadly necessary support for scientifically sound restoration. EPIL-connected institutes, agencies, and individual experts have varying capacities, research mandates, and administrative interests, resulting in varying levels of knowledge and experience in ecological science. Nevertheless, their contributions to the appraisal process are crucial for court judgments that determine fish release practices. The current judicial practices, combined with a lack of biological and environmental expertise in local courts, have resulted in potentially significant environmental and biological challenges.

### Off-target ecological conservation efforts

Species of economic significance are often prioritized by the judicial systems in fish conservation in the Yangtze River basin (Fig. 3). However, effective conservation relies on maintaining a healthy ecosystem, not just artificially increasing the populations of targeted species. Many non-CAS and non-VES play crucial roles in preserving ecosystem stability and sustainability. For example, *Hemiculter leucisculus*, a non-CAS, serves as an important food source for endangered species, such as *Neophocaena asiaeorientalis* (Nabi et al. 2017, Yang et al. 2019). Similarly, species in the genus *Rhodeus* maintain symbiotic relationships with freshwater mussels and are crucial for mussel reproduction (Kitamura et al. 2005, Yu et al. 2021). Both *Rhodeus spp. and Hemiculter leucisculus* play significant roles in sustaining a healthy food web, supporting water quality, and facilitating energy transfer in the Yangtze River ecosystem (Li et al. 2021, Guo et al. 2022a,b). However, these taxa were represented in 11 court cases as being captured without being released, likely due to their low value as a fisheries resource.

The mismatches between the taxa of fishes caught and those released have potentially reinforced the basin-wide prevalence of CAS, as these species are more likely to occupy the entire river system compared to other species, particularly VES. Among the 21 species in the caught-and-released group, 16 were whole-river-occupying species and five were species endemic to the upstream and midstream sections of the basin (Fig. 4). In contrast, of the 22 taxa in the caught-but-not-released group, only eight were whole-river-occupying, while the remaining 14 species were endemic to specific parts of Yangtze River basin. The weakening of locally endemic species and strengthening of whole-river-occupying species suggests a potential detrimental trend towards the homogenization of the riverine fauna across the basin (Rahel 2002).

**Figure 4.**
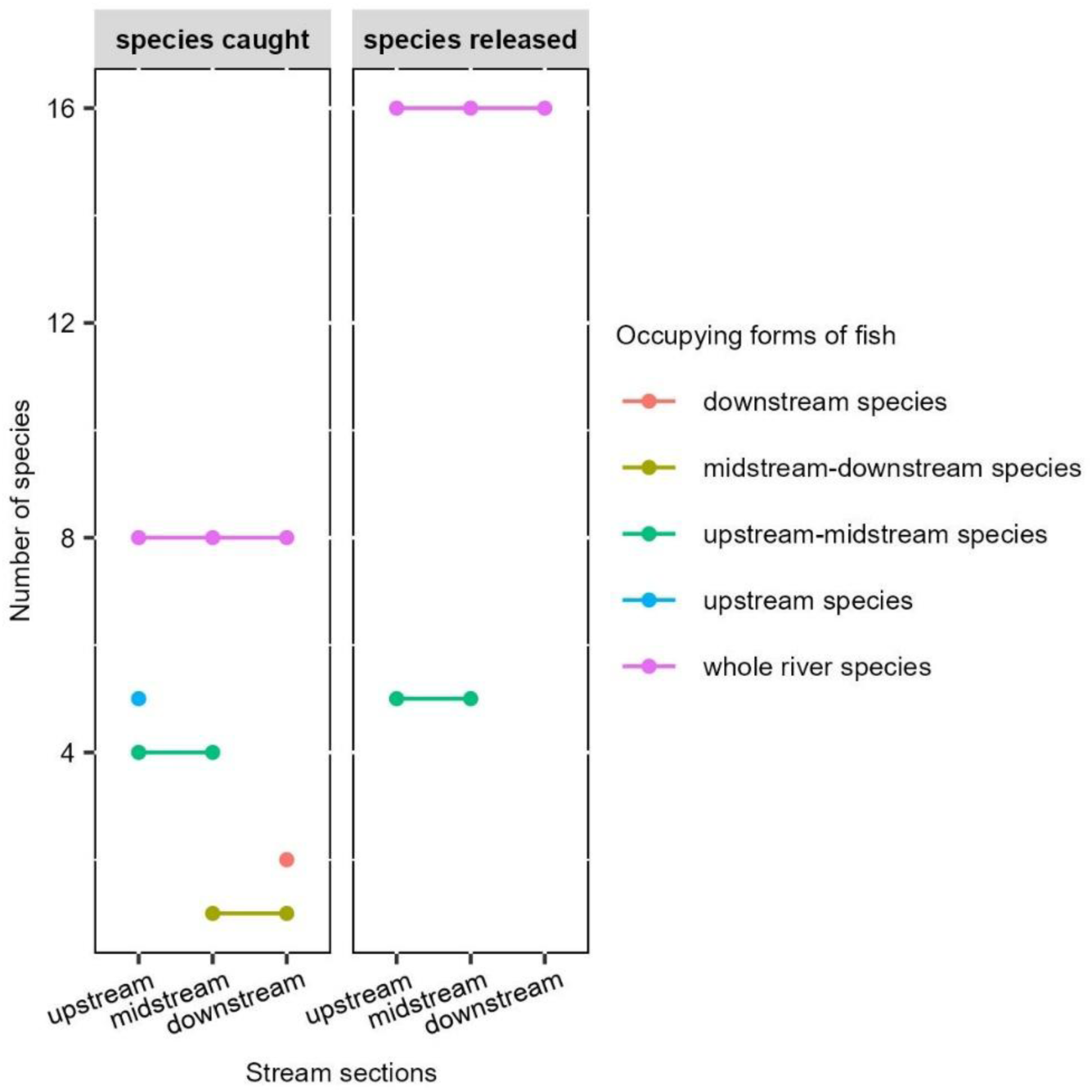
Distribution of fish species by stream section in illegal fishing cases along the Yangtze River. Fish species involved in illegal fishing cases are categorized based on their geographic distribution within the Yangtze River: upstream, midstream, downstream, or across multiple sections. Individual points represent species restricted to a single river section, while connecting lines indicate species distributed across two or more sections.

Inaccurate taxonomic identification can misguide conservation efforts and carry significant ecological consequences. Such errors may stem from a lack of scientific expertise within the judicial system, resulting in species misidentification (e.g., confusion between *Coilia brachygnathus* and *Coilia mystus*), disregard for the ecological roles of certain species (e.g., *Rhodeus spp.*), or grouping of species with similar commercial value (e.g., *Culterinae spp.*). These inaccuracies impede an accurate assessment of the Yangtze River’s ecological condition and compromise the effective protection of the target species. For example, identifying *Rhodeus* spp. only to the genus level overlooks species-specific conservation needs, as this genus comprises five species with varying levels of threatened status in the Yangtze River basin (Yang et al. 2023). Moreover, misidentification during court-ordered releases can result in inappropriate substitutions based on superficial similarities, such as foraging strategies, which may intensify ecological competition (Fig. 5, Table S4). For example, when the benthopelagic carnivore *Culter alburnus* is substituted with *Chanodichthys mongolicus mongolicus*, another benthopelagic carnivore, the competition between these two species could undermine the restoration efforts for *Culter alburnus*. In 11 of the 12 substitution cases reviewed, the foraging strategies of the released taxa differed from those of the captured taxa (Fig. 5, Table S4), potentially disrupting food webs and energy flows and exacerbating the ecological imbalances caused by illegal fishing. In one extreme example, illegally captured shrimp and crab were replaced with *Hypophthalmichthys molitrix*, an species from an entirely different phylum, with no consideration for the distinct ecological roles played by each organism.

**Figure 5.**
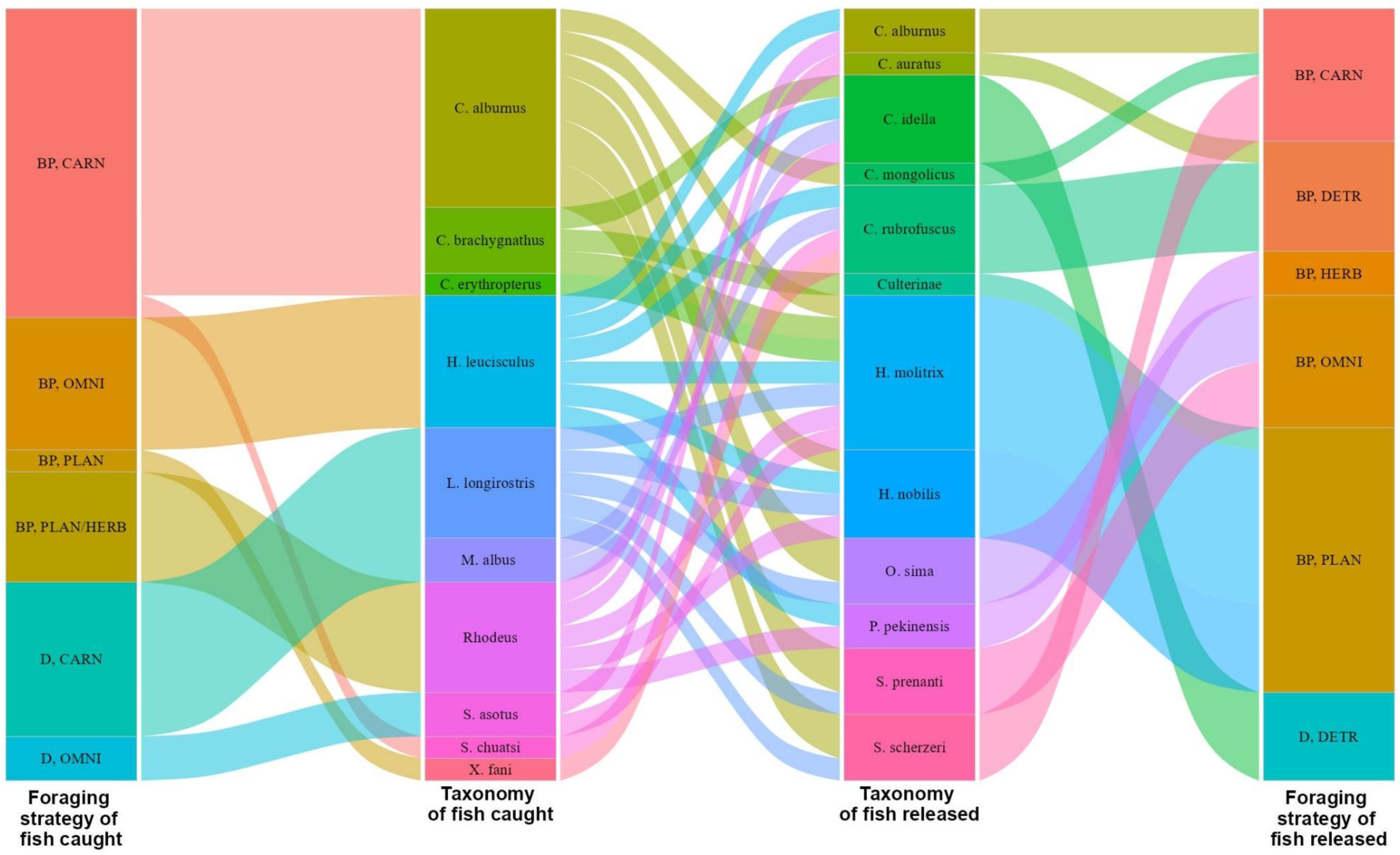
Fish substitution patterns by foraging strategy and habitat in all the mismatch cases (n=18) with known taxonomic names of both illegally caught and court-ordered released taxa. Strata in the third and fourth columns represent the times fish from each category were used as substitutes. The width of each flow between caught and released taxa is proportional to the number of instances in which a caught taxon was substituted by a different taxon as court-ordered. Abbreviations: BP = benthopelagic; D = demersal; CARN = carnivores; DETR = detritivores; HERB = herbivores; OMNI = omnivores; PLAN = planktivores.

Genetic contamination of natural populations caused by the release of artificially raised fish has already impacted wild populations of *Hypophthalmichthys molitrix* (Fang et al. 2021), *Carassius auratus* (Feng et al 2023), and *Procypris rabaudi* (Cheng et al. 2011) in the Yangtze River basin, all before and during the implementation of the Environmental Protection and Information Law (EPIL). *Hypophthalmichthys molitrix* was released as a remedy in 24 illegal fishing cases, making it the second most frequently released species, while *Procypris rabaudi* was released despite not being among the species caught illegally. In addition to concerns about genetic pollution, habitat degradation due to nutrient loading resulting from the 75% mortality rate of the released fish is also a potential concern. Ironically, being aware of the high post-release mortality, judges ordered the release of biomass up to four times that of the captured fish—and sometimes an additional fourfold of the estimated offspring from eggs the captured fish may have carried—reaching several thousand individuals in some cases (Table S2), in order to meet compensation guidelines, without considering the broader ecological consequences of such actions.

Environmental protection laws have been implemented globally, yet their effectiveness is often under-evaluated. Overall, illegal fishing EPIL cases increased in the Yangtze River basin from 2015 to 2019, reflecting the growing enforcement efforts of public prosecutors in public-interest litigation. However, the lack of scientific expertise in judicial remedies has led to frequent misidentification of fish taxa, mismatches between the species caught and released, and a bias toward economically valuable species. In this study, we highlight the disconnect between the environmental protection goals and the actual ecological outcomes of the Yangtze River’s judicial system. These discrepancies not only undermine ecosystem sustainability but may actively hinder it. At the core of this problem is a failure to integrate ecological protection knowledge into the judicial process, particularly in decisions regarding restoration remedies. Compounding this issue is a prevailing bias that prioritizes the economic value of natural resources. While economic considerations are important, relying on them exclusively can lead to damaging outcomes, especially when ecological factors such as population recovery, habitat integrity, genetic diversity, and interspecies dynamics are overlooked. These challenges reveal a deeper disconnect between social and ecological disciplines, underscoring the urgent need for interdisciplinary collaboration to build judicial systems capable of supporting both social and environmental sustainability.

## Research Methods

We conducted a case-based analysis of Environmental Public Interest Litigation (EPIL) in China, focusing on the Yangtze River basin between 2015 and 2021. First, we reviewed the national development of EPIL policy during this period to provide contextual background. We then systematically examined court decisions related to EPIL cases in the Yangtze River region, with particular attention to the types of environmental restoration remedies ordered by judges for illegal fishing cases. Our analysis assessed both the accuracy of these remedies in addressing the ecological harms identified in each case and their potential effectiveness in contributing to environmental or ecosystem recovery.

### Collection of court files

We employed a legal-empirical research approach to conduct a textual analysis of court rulings related to Environmental Public Interest Litigation (EPIL). Court documents from 2015 to 2021 were obtained primarily from China Judgment Online (https://wenshu.court.gov.cn/website/wenshu/181029CR4M5A62CH/index.html), the official legal document database maintained by the Supreme People’s Court of China. Additional case records were cross-referenced using Peking Law (https://www.pkulaw.com/) and OpenLaw (http://www.openlaw.cn) databases to ensure completeness. The following keywords were used to retrieve relevant cases: “environmental public interest litigation,” “Yangtze River,” “Sichuan Province,” “Hubei Province,” and “Jiangsu Province.” These three provinces were selected to represent the upper, middle, and lower reaches of the Yangtze River Basin, respectively (Fig. 2), encompassing distinct ecological and geographical characteristics. Collectively, the mainstream and tributaries passing through these provinces support over 95% of the basin’s fish species (Fu et al., 2003), with notable variation in community composition (Kang et al., 2018). In addition, these provinces contribute substantially to China’s freshwater fisheries economy (Table S).

The initial court file screening process identified 516 EPIL cases for further analysis (Fig. 2). These cases encompassed a wide range of environmental violations, including but not limited to illegal fishing, water pollution, deforestation, unauthorized construction, illegal mining, poaching, noise pollution, improper use of agricultural land, waste gas emissions, and illegal disposal of solid waste. From this dataset, we narrowed our focus to 120 cases specifically involving illegal fishing (Fig. 1b–d), due to their prevalence and direct ecological relevance to the Yangtze River system. Each of these cases was examined in detail to assess the judicial decision-making process, the types of environmental restoration remedies ordered, and the implications of these remedies for ecosystem sustainability in the Yangtze River Basin.

### Extractioin of Fish-Related Information from Court Files

From the 120 illegal fishing cases, we extracted detailed information on the fish involved in each case. This included data on both the remedial methods implemented and the taxa of fish that were either caught or released as part of court-ordered restoration efforts. Specifically, we recorded the taxonomic identity and abundance (measured as biomass or number of individuals) of each fish taxon, as well as their habitate and foraging strategy types and the geographic locations where the cases occurred. To ensure the accuracy of species identification, we cross-validated the reported taxa against their known geographic distributions using authoritative references (Ding, 1994; Fu et al., 2003; Yang et al., 2023).

### Definition of fish groups used in this research

To illustrate the taxonomic composition of fish involved in the illegal fishing cases and to interpret their presence in court-ordered remedies, we categorized all recorded taxa based on their ecological status and economic importance. Vulnerable and Endangered Species (VES) include fish classified as "Vulnerable," "Endangered," or "Critically Endangered" according to the latest IUCN Red List assessments (2024). Common Aquaculture Species (CAS) refer to the 14 most widely produced aquaculture species across the three focal provinces (Table S5). These include the four major Chinese carps (*Mylopharyngodon piceus*, *Hypophthalmichthys molitrix*, *Hypophthalmichthys nobilis*, and *Ctenopharyngodon idella*), as well as ten other commonly farmed species (*Carassius auratus*, *Cyprinus rubrofuscus*, *Parabramis pekinensis*, *Tachysurus fulvidraco*, *Monopterus albus*, *Silurus asotus*, *Siniperca chuatsi*, *Misgurnus anguillicaudatus*, and *Channa argus*). Together, these species account for approximately 7,094,608 (±198,135.14) metric tons—96.4% of the total annual aquaculture production—according to the China Fishery Statistics Yearbook (Table S5).

### Comparison of Fish Species Illegally Caught and Court-Ordered for Release

Based on case records, fish taxa were classified into three groups: (1) species both illegally caught and court-ordered for release, (2) species illegally caught but not released, and (3) species released by court order but not originally caught (Table 1). We compared taxonomic differences among these groups depending on data availability. For all fish taxa with identifiable names, we quantified their frequency of occurrence across the three categories. In cases where both the caught and released taxa were named, we examined potential species substitution and compared the ecological status (e.g., IUCN Red List category) and economic importance (e.g., annual aquaculture production) of the substituted species (Figs. 3 & 4).

For the Common Aquaculture Species (CAS), we used linear regression models to examine the relationship between provincial annual aquaculture production (measured in metric tons) and the number of instances in which each species was (a) illegally caught or (b) court-ordered for release. These models were fitted using a negative binomial distribution (glm.nb) to account for overdispersion in count data. Diagnostic results of the *glm.nb* models indicated a good fit to the data (Extended Fig. 2–4). We then used paired bootstrap resampling (1,000 iterations) to compare the strength of the two relationships—between production and illegal catch, and between production and court-ordered release—by estimating the distribution of slope differences (Extended Fig. 1). Statistical significance was determined using a two-sided bootstrap p-value, with α = 0.05 as the threshold for significance.

## Supplementary information

**Table S1.**
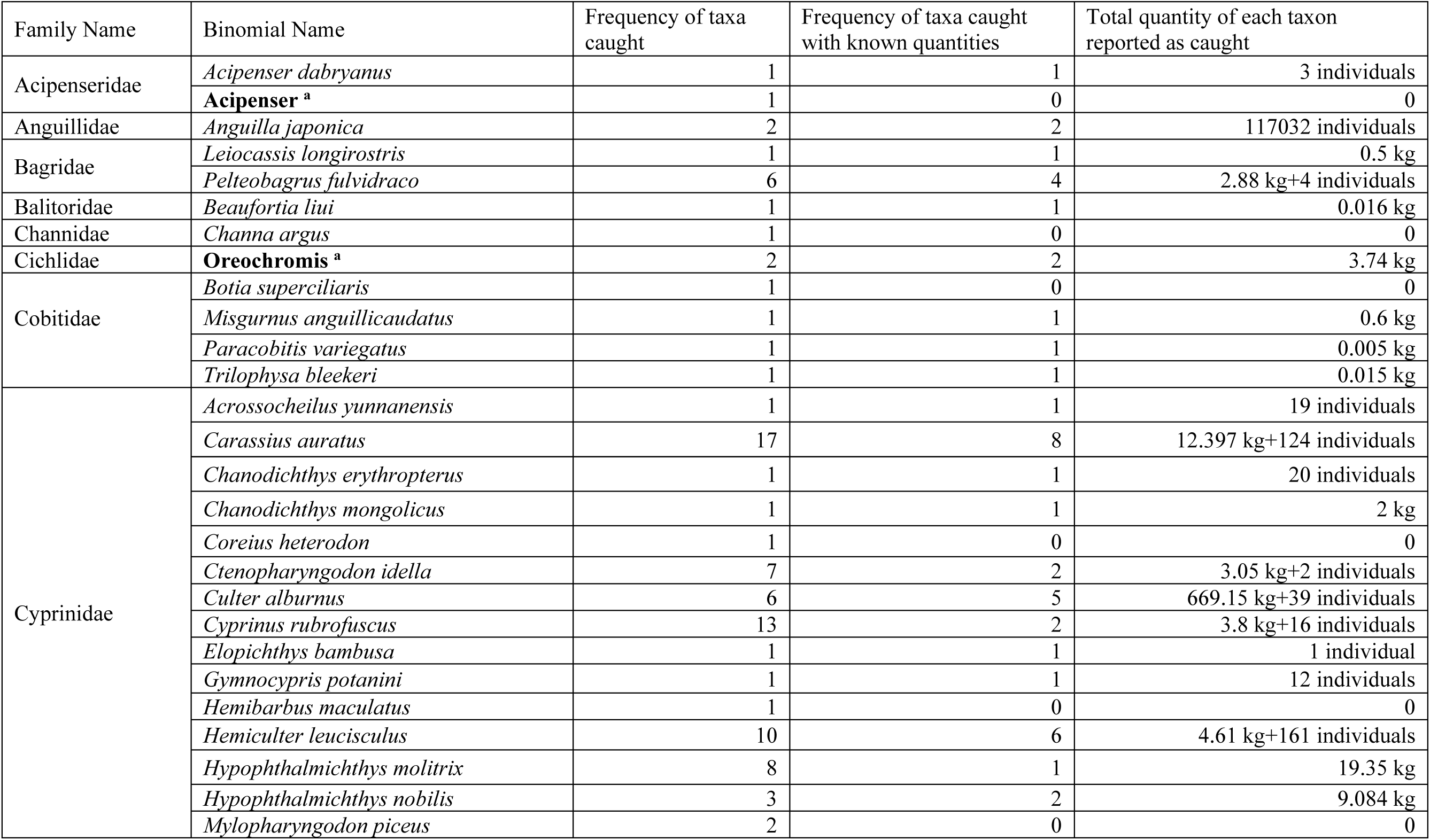
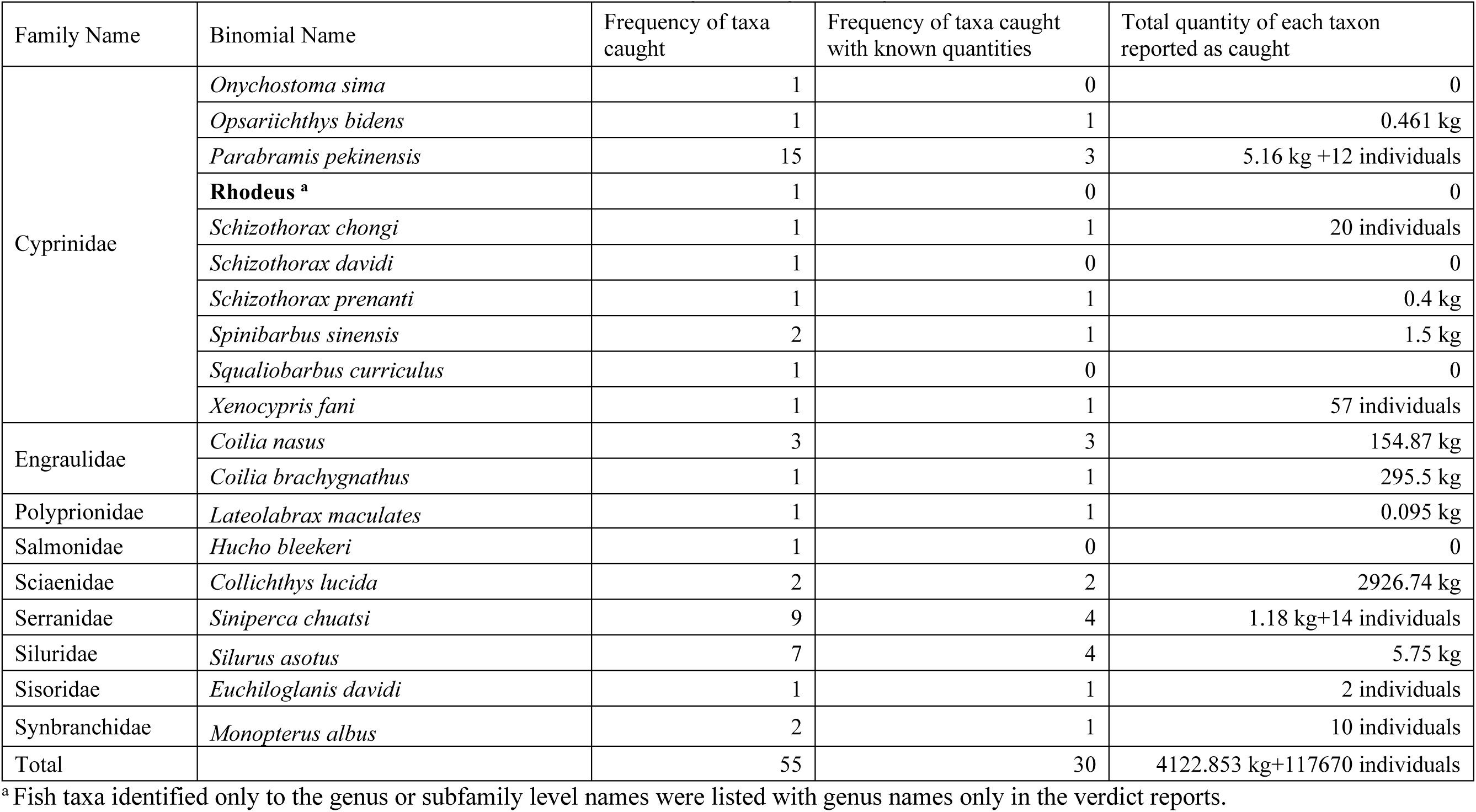
Biomass and instance of each fish taxon caught in illegal fishing cases.

**Table S2.** Biomass and instance of each fish taxon court ordered-released in illegal fishing cases.

| Family Name | Binomial Name | Frequency of release | Frequency of release with known quantities | Number of juveniles released | Weight of adult fish released (kg) |
| --- | --- | --- | --- | --- | --- |
| Bagridae | *Leiocassis longirostris* | 1 | 1 | 98 | 1 |
|  | *Procypris rabaudi* | 1 | 1 | 100 | / |
| Channidae | *Channa argus* | 4 | 3 | 6700 | 12.7 |
| Cobitidae | *Leptobotia elongata* | 1 | 1 | 100 | / |
|  | *Misgurnus anguillicaudatus* | 2 | 2 | 5000 | 0.46 |
| Cyprinidae | *Carassius auratus* | 32 | 31 | 342790 | 304.78 |
|  | *Chanodichthys mongolicus* | 7 | 7 | 12450 | 8.7 |
|  | *Ctenopharyngodon idella* | 20 | 19 | 254569 | 233.4 |
|  | *Culter alburnus* | 2 | 1 | 1000 | 1.081 |
|  | **Culterinae<sup>a</sup>** | 6 | 6 | 324840 | 511.6 |
|  | *Cyprinus rubrofasciatus* | 24 | 23 | 413431 | 435.58 |
|  | *Hypophthalmichthys molitrix* | 24 | 23 | 580635 | 106.8 |
|  | *Hypophthalmichthys nobilis* | 15 | 15 | 238966 | 61.15 |
|  | *Mylopharyngodon piceus* | 3 | 3 | 12000 | / |
|  | *Onychostoma sima* | 4 | 4 | 1300 | / |
|  | *Parabramis pekinensis* | 8 | 6 | 122800 | 363 |
|  | *Schizothorax davidi* | 1 | 1 | 1800 | / |
|  | *Schizothorax prenanti* | 2 | 2 | 2000 | / |
|  | **Schizothorax<sup>a</sup>** | 1 | 1 | 300 | / |
|  | *Spinibarbus sinensis* | 1 | 1 | 10000 | 17.44 |
|  | *Pelteobagrus fulvidraco* | 8 | 7 | 32769 | 13.26 |
| Serranidae | *Siniperca scherzeri* | 3 | 3 | 1500 | / |
|  | *Siniperca chuatsi* | 11 | 11 | 19681 | 35.64 |
| Siluridae | *Silurus asotus Linnaeus* | 4 | 4 | 5670 | 11.3 |
| Synbranchidae | *Monopterus albus* | 1 | / | / | 2.47 |
| Total |  | 49 | 44 | 2390499 | 2120.361 |

<sup>a</sup> Fish taxa identified only to the genus or subfamily level names were listed with genus names only in the verdict reports.

**Table S3.** Frequency of fish and aquatic animal taxa in cases where released taxa differed from those illegally caught.

| Taxa | Frequency of taxa | Frequency of release for substitution | Frequency of taxa released but not caught | Frequency of taxa caught but not released | Frequency of taxa substituted by other taxa |
| --- | --- | --- | --- | --- | --- |
| *Botia supercilialis* | 1 |  |  | 1 |  |
| *Carassius auratus*<sup>a</sup> | 2 | 2 |  |  |  |
| *Chanodichthys mongolicus* | 1 | 1 |  |  |  |
| *Coilia brachygnathus* | 1 |  |  |  | 1 |
| *Ctenopharyngodon idella*<sup>a</sup> | 4 | 3 | 1 |  |  |
| *Culter alburnus* | 4 | 1 |  |  | 3 |
| **Culterinae**<sup>b</sup> | 2 | 1 | 1 |  |  |
| *Cultrichthys erythropterus* | 1 |  |  |  | 1 |
| *Cyprinus rubrofasciatus*<sup>a</sup> | 3 | 3 |  |  |  |
| *Hemiculter leucisculus* | 3 |  |  | 1 | 2 |
| *Hucho bleekeri* | 1 |  |  | 1 |  |
| *Hypophthalmichthys molitrix*<sup>a</sup> | 7 | 6 | 1 |  |  |
| *Hypophthalmichthys nobilis*<sup>a</sup> | 4 | 2 | 2 |  |  |
| *Leiocassis longirostris* | 1 |  |  |  | 1 |
| *Monopterus albus* | 2 |  |  | 1 | 1 |
| *Mylopharyngodon piceus*<sup>a</sup> | 1 |  | 1 |  |  |
| *Onychostoma sima* | 3 | 2 | 1 |  |  |
| *Parabramis pekinensis*<sup>a</sup> | 1 | 1 |  |  |  |
| **Rhodeus**<sup>b</sup> | 1 |  |  |  | 1 |
| *Schizothorax prenanti* | 2 | 2 |  |  |  |
| *Silurus asotus* | 1 |  |  |  | 1 |
| *Siniperca chuatsi* | 1 |  |  |  | 1 |
| *Siniperca scherzeri* | 3 | 2 | 1 |  |  |
| *Xenocypris fani* | 1 |  |  |  | 1 |
| Shrimp | 2 |  |  |  | 2 |
| Crab | 1 |  |  |  | 1 |
| Turtle | 1 |  |  |  | 1 |

<sup>a</sup> Common Aquaculture Species (CAS) based on the Chinese Fishery statistics Yearbook (2016-2022).
<sup>b</sup> Fish taxa identified only to the genus or subfamily level names were listed with genus names only in the verdict reports.

**Table S4.**
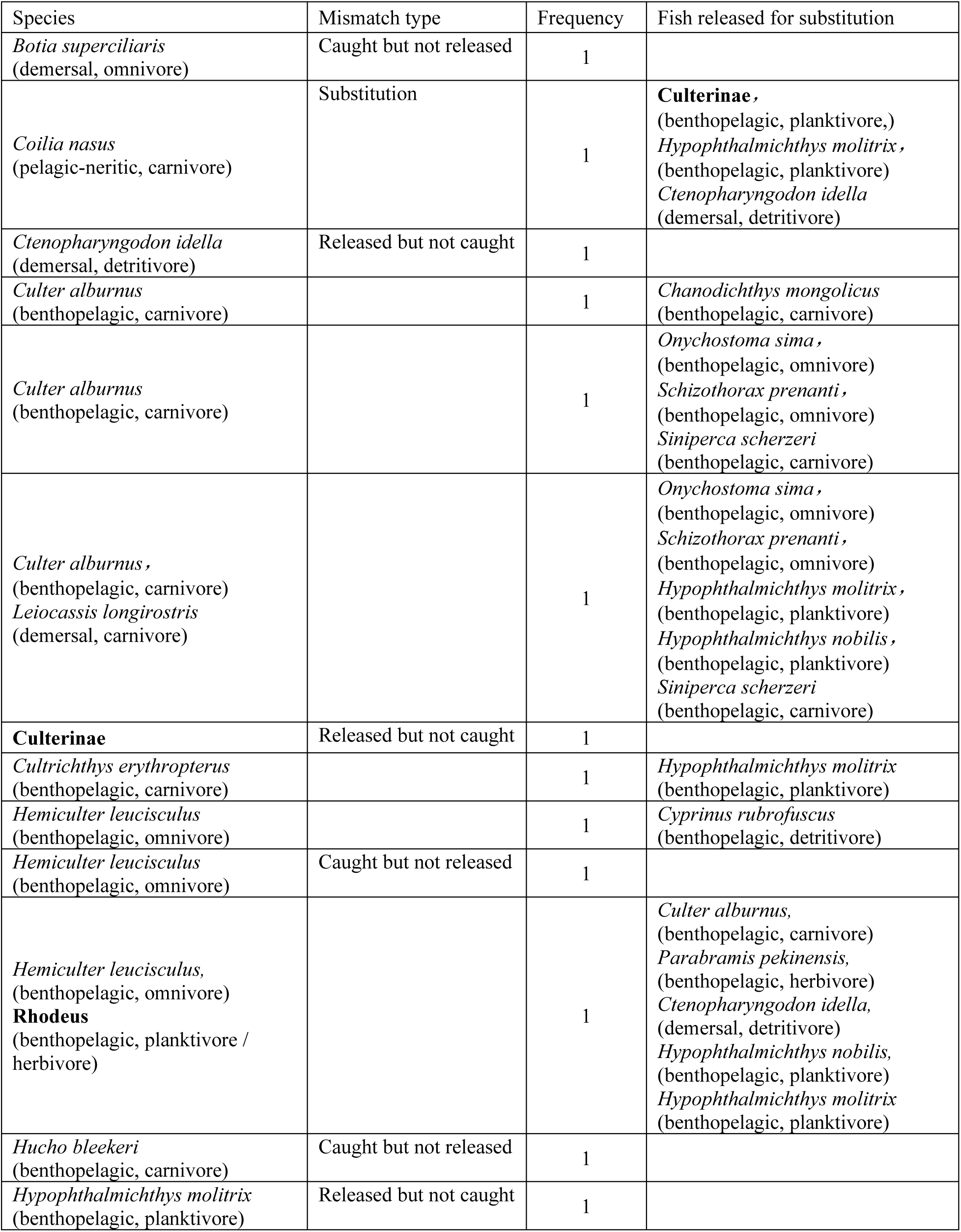
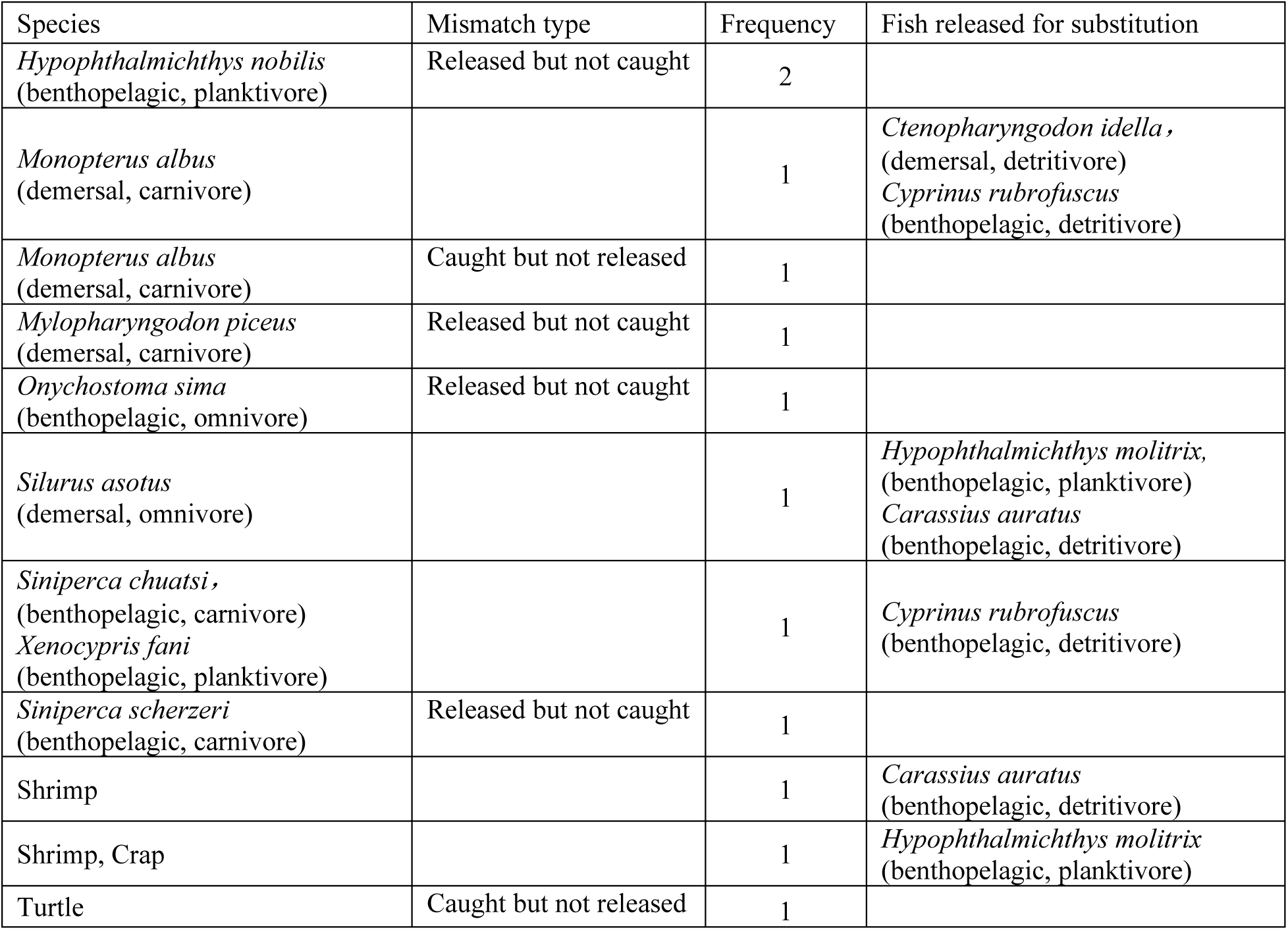
Foraging strategies and habitat types of taxa involved in mismatches. Foraging types of the two subfamilies Culterinae and Rhodeus were based on likely species at the locations where the cases occurred.

**Table S5.** Average annual production of Common Aquaculture Species in the three provinces from 2015 to 2021. Data were extracted from the *China Fishery Statistics Yearbooks* (2016–2022). Values in parentheses represent the standard deviation of annual production during this period.

| Province | Sichuan |  | Hubei |  | Jiangsu |  | Sum of three provinces |  |
| --- | --- | --- | --- | --- | --- | --- | --- | --- |
|  | Weight<br>(metric ton) | Portion | Weight<br>(metric ton) | Portion | Weight<br>(metric ton) | Portion | Weight<br>(metric ton) | Portion |
| Sum of all species | 1457124.71<br>(±92352.90) | 1.00 | 3488156.57 (±<br>150001.28) | 1.00 | 2415053.86(±<br>100855.17) | 1.00 | 7122167(±<br>198135.14) | 1.00 |
| *Ctenopharyngodon idella* | 265785.57(±<br>22588.16) | 0.18 | 906523.14(±<br>37722.68) | 0.26 | 418833.71(±<br>23716.18) | 0.17 | 1591142.43(±<br>48928.93) | 0.22 |
| *Hypophthalmichthys molitrix* | 308260.14(±<br>13899.87) | 0.21 | 602197.57(±<br>79603.33) | 0.17 | 447526.57(±<br>29807.81) | 0.19 | 1357984.29(±<br>95454.72) | 0.18 |
| *Carassius auratus* | 189943.29(±<br>16469.06) | 0.13 | 400280.29(±<br>62256.43) | 0.11 | 623959.86(±<br>15896.22) | 0.26 | 1214183.43(±<br>56820.26) | 0.16 |
| *Hypophthalmichthys nobilis* | 171322.86(±<br>12532.73) | 0.12 | 445337.14(±<br>14579.91) | 0.13 | 243209.86(±<br>13619.68) | 0.10 | 859869.86(±<br>29950.41) | 0.12 |
| *Cyprinus rubrofuscus* | 184677.29(±<br>10210.56) | 0.13 | 144437.71(±<br>38052.15) | 0.04 | 147277.86(±<br>8636.73) | 0.06 | 476392.86(±<br>33787.36) | 0.06 |
| *Parabramis pekinensis* | 32803.14(±<br>1840.57) | 0.02 | 232104.14(±<br>19575.71) | 0.07 | 170835.00(±<br>10315.08) | 0.07 | 435742.29(±<br>15515.21) | 0.06 |
| *Mylopharyngodon piceus* | 2053.86(±<br>129.83) | 0.00 | 177954.57(±<br>41183.16) | 0.05 | 92815.14(±<br>5316.09) | 0.04 | 272823.57(±<br>42336.98) | 0.04 |
| *Tachysurus fulvidraco* | 31921.57(±<br>3893.07) | 0.02 | 126649.57(±<br>21698.89) | 0.04 | 26064.00(±<br>1617.60) | 0.01 | 184635.14(±<br>25061.31) | 0.03 |
| *Monopterus albus* | 11226.29(±<br>709.55) | 0.01 | 156008.14(±<br>19891.39) | 0.04 | 4561.29(±<br>1956.79) | 0.00 | 171795.71(±<br>21605.60) | 0.02 |
| *Silurus asdus* | 86597.86(±<br>12673.54) | 0.06 | 46116.00(±<br>14388.30) | 0.01 | 1142.86(±<br>419.38) | 0.00 | 133856.71(±<br>24868.33) | 0.02 |
| *Siniperca chuatsi* | 2463.57(±<br>393.58) | 0.00 | 66266.57(±<br>16459.18) | 0.02 | 26518.57(±<br>6957.70) | 0.01 | 95248.71(±<br>18008.26) | 0.01 |
| *Misgurnus anguillicaudatus* | 31449.43(±<br>2332.02) | 0.02 | 41982.86(±<br>4597.50) | 0.01 | 48829.29(±<br>8805.26) | 0.02 | 122261.57(±<br>10834.35) | 0.02 |
| *Silurus asdus* | 79013.29(±<br>4172.64) | 0.05 | 24375.86(±<br>11278.24) | 0.01 | 6510.86(±<br>2160.70) | 0.00 | 109900.00(±<br>16286.95) | 0.01 |
| *Channa argus* | 9740.71(±<br>431.61) | 0.01 | 31788.14(±<br>7683.74) | 0.01 | 27242.71(±<br>3239.70) | 0.01 | 68771.57(±<br>10402.56) | 0.01 |
| Others | 49865.86(±<br>6653.31) | 0.03 | 86134.86(±<br>33033.70) | 0.02 | 129726.29(±<br>24750.56) | 0.05 | 265727.00(±<br>54380.25) | 0.04 |

## Extended Figures

**Extended Figure 1.**
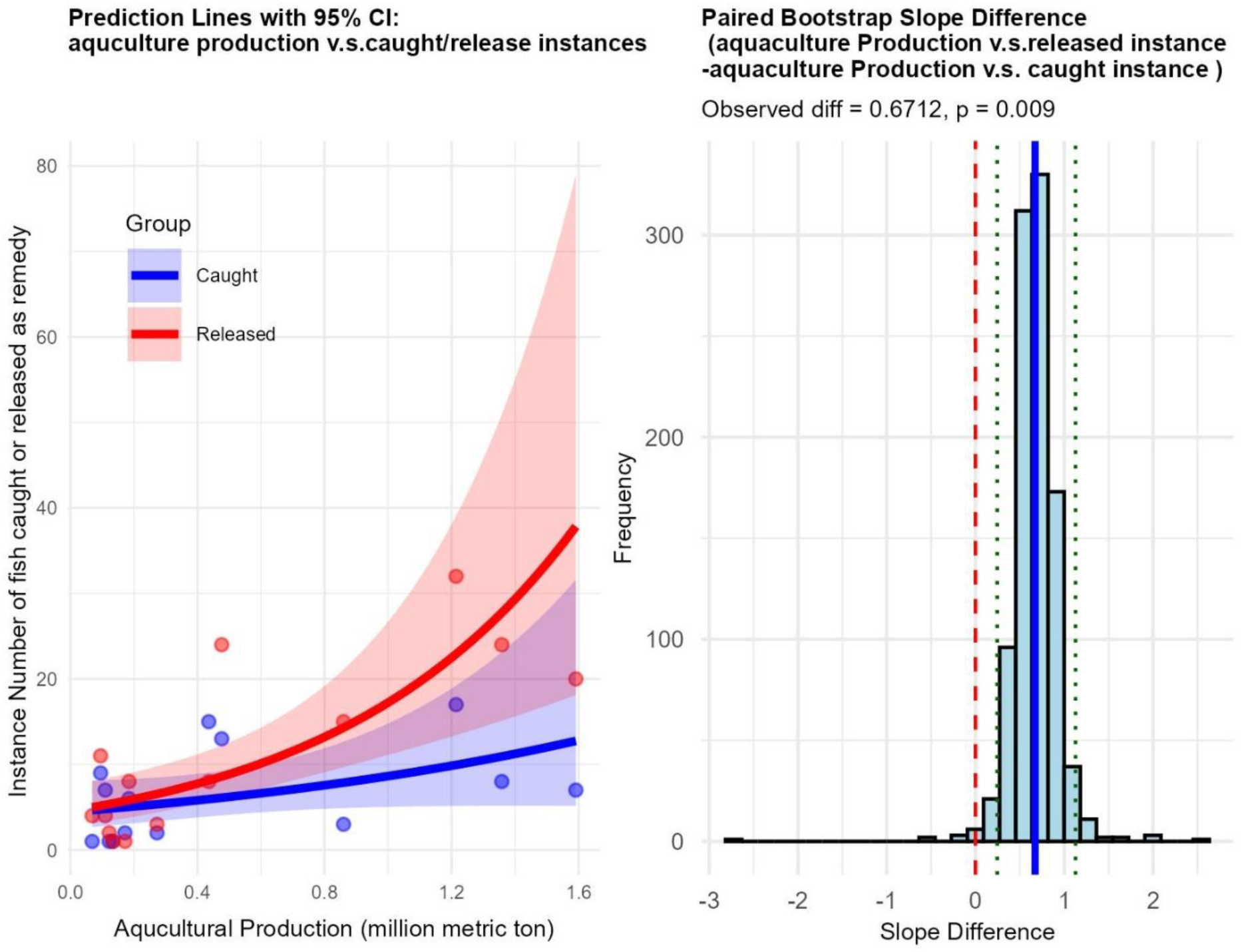
Comparison of slope estimates from two negative binomial regression models relating annual aquatic production to (1) illegal fishing instances and (2) court-ordered release instances of 14 commonly aquacultured species in three Yangtze River provinces. (a) Slope estimates with 95% confidence intervals for each model. (b) Paired bootstrap distribution of the slope difference between the two models.

**Extended Figure 2.**
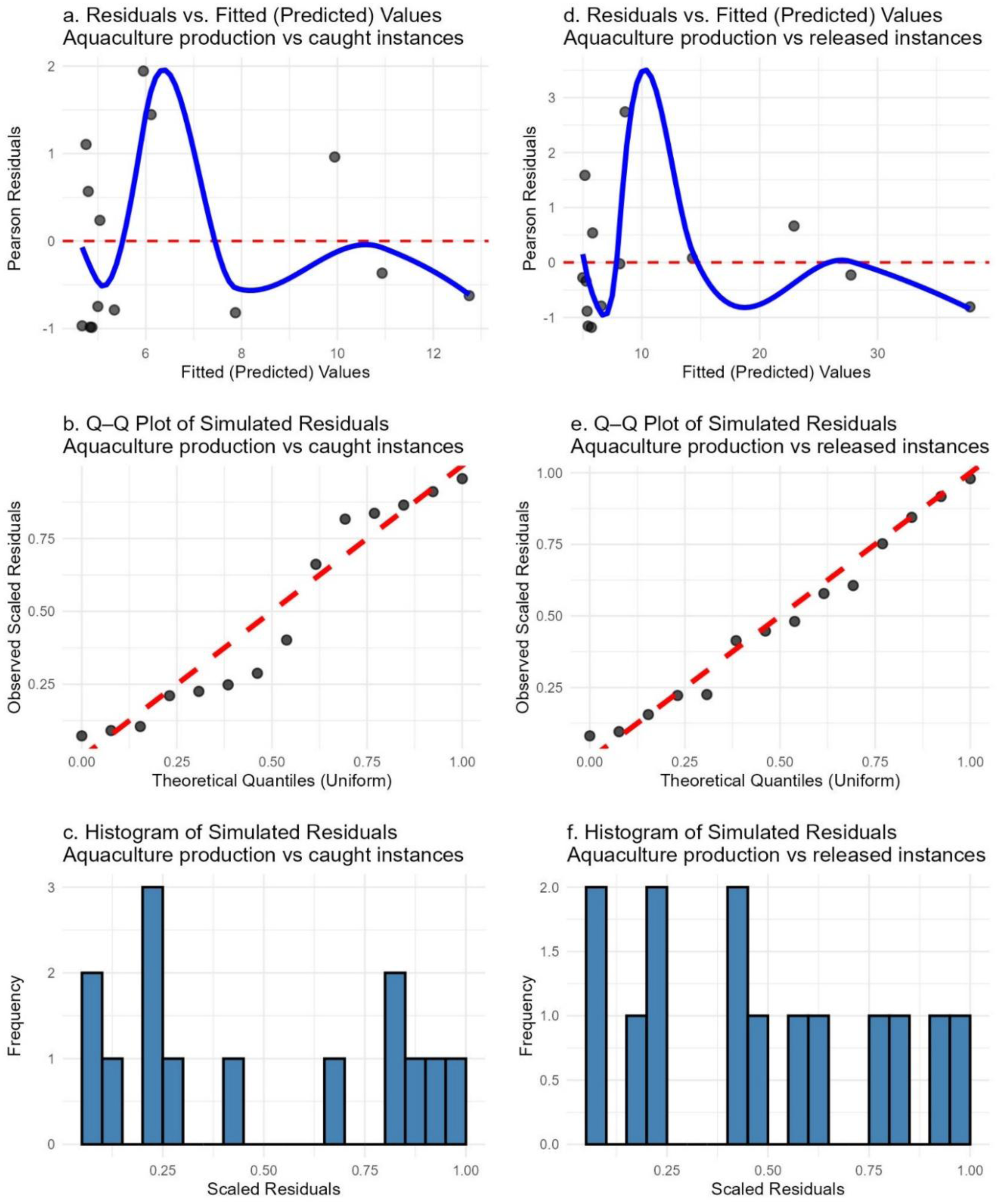
Diagnostic plots for two negative binomial regression models analyzing relationships between annual aquatic production and enforcement-related activities involving 14 commonly aquacultured species in three provinces along the Yangtze River. Panels (a–c) correspond to the model that relates annual aquatic production to the number of illegal fishing instances; panels (d–f) correspond to the model that relates annual aquatic production to the number of court-ordered release instances. (a, d) Residuals vs. fitted values assessing linearity and homoscedasticity; (b, e) Q–Q plots of simulated residuals evaluating deviations from normality; and (c, f) Histograms of simulated residuals examining residual distributions. These diagnostics indicate that both models provided a good fit to the data.

**Extended Figure 3.**
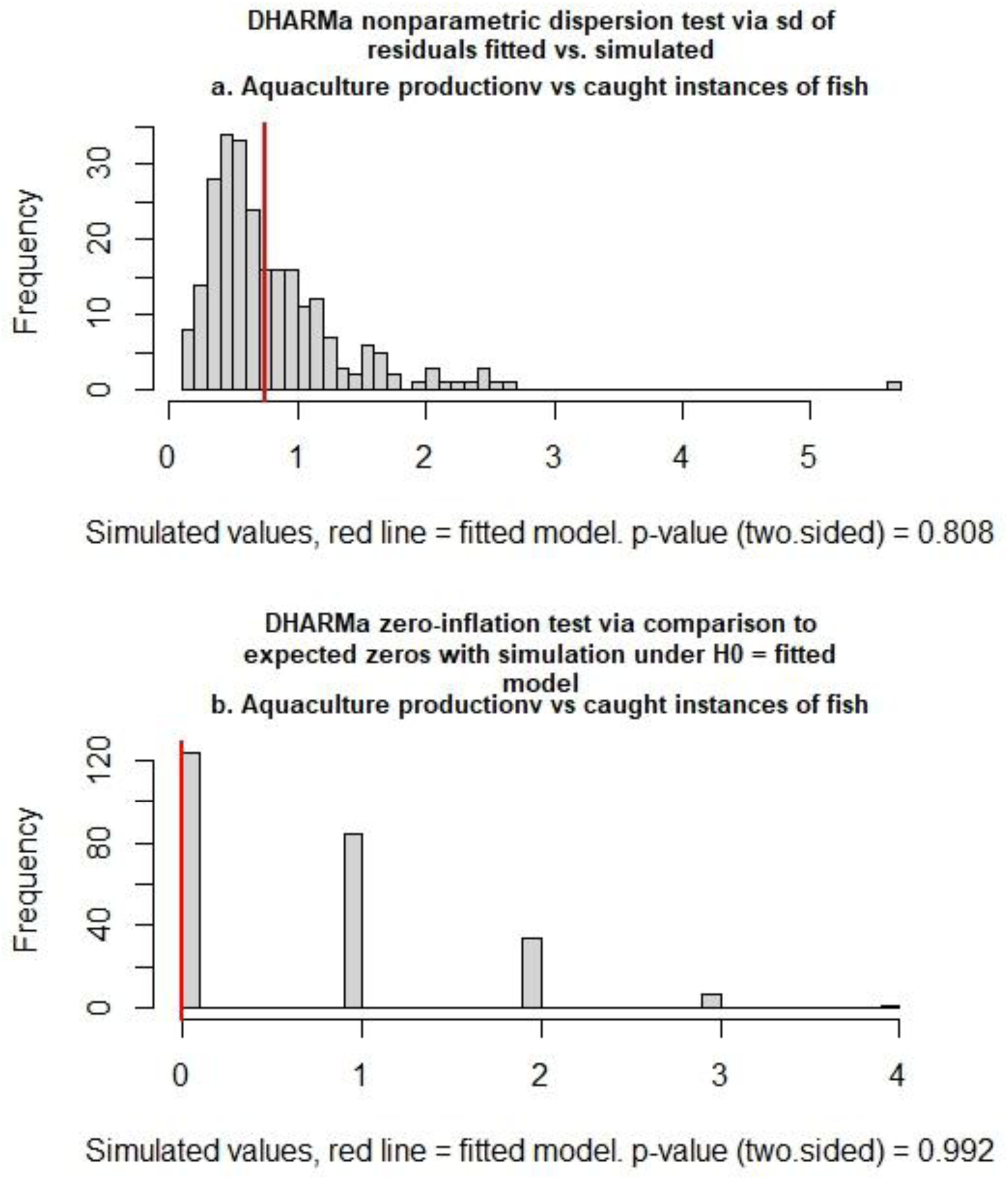
DHARMa diagnostic tests for the negative binomial regression models relating annual aquatic production to illegal fishing instances of 14 aquacultured species in three provinces along the Yangtze River. a. Non-parametric dispersion test assessing residual overdispersion. b. Zero-inflation test comparing observed and expected zeros under the fitted model (H₀). No significant zero-inflation was detected.

**Extended Figure 4.**
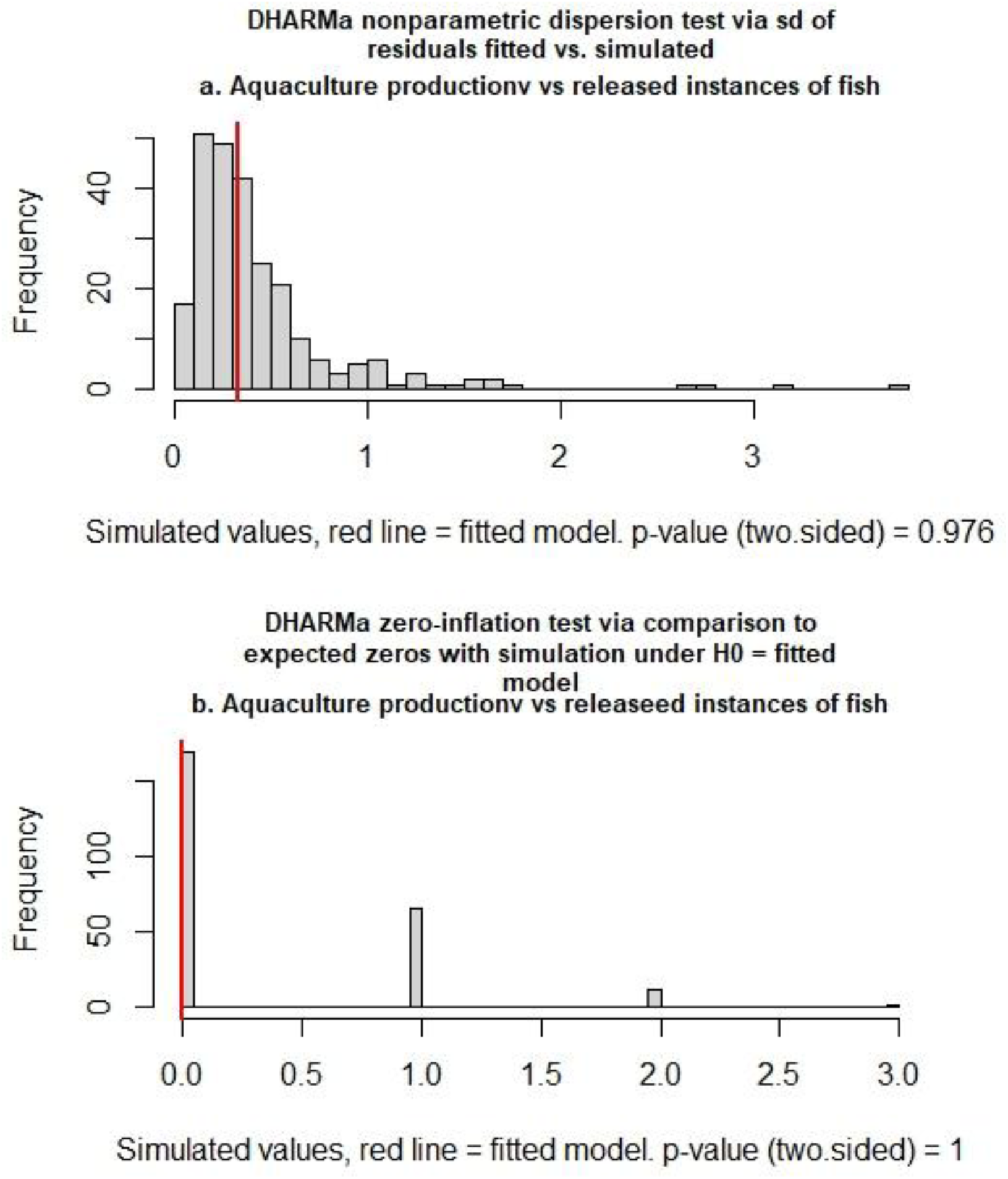
DHARMa diagnostic tests for the negative binomial regression models relating annual aquatic production to court-ordered release instances of 14 aquacultured species in three provinces along the Yangtze River. a. Non-parametric dispersion test assessing residual overdispersion. b. Zero-inflation test comparing observed and expected zeros under the fitted model (H₀). No significant zero-inflation was detected.

## References

Bell, S., McGillivray, D. & Pedersen, O. Environmental Law Ch. 1, p. 4 (Oxford Univ. Press, 2013).

Chen, D. et al. Status of research on Yangtze fish biology and fisheries. Environ. Biol. Fishes 85, 337–357 (2009).

Chen, T., Wang, Y., Gardner, C. & Wu, F. Threats and protection policies of the aquatic biodiversity in the Yangtze River. *J*. Nat. Conserv. 58, 125931 (2020).

Cheng, F. et al. Impacts of hatchery release on genetic structure of rock carp *Procypris rabaudi* in the upper Yangtze River, China. Fish. Sci. 77, 765–771 (2011).

Deng, Y., Zhao, J., Lu, G., Wu, X. & Tao, Y. Cloning, characterization and expression of the pepsinogen C from the golden mandarin fish *Siniperca scherzeri* (Teleostei: Perciformes). Fish. Sci. 76, 819–826 (2010).

Ding, R. The Fish of Sichuan (Sichuan Science and Technology Press, 1994).

Fang, D. et al. Relationship between genetic risk and stock enhancement of the silver carp (*Hypophthalmichthys molitrix*) in the Yangtze River. Fish. Res. 235, 105829 (2021).

Feng, X. et al. Genetic responses in sexual diploid and unisexual triploid goldfish (*Carassius auratus*) introduced into a high-altitude environment. Mol. Ecol. 32, 1955–1971 (2023).

Food and Agriculture Organization of the United Nations. The state of world fisheries and aquaculture 2016. FAO, Rome (2016).

Fu, C., Wu, J., Chen, J., Wu, Q. & Lei, G. Freshwater fish biodiversity in the Yangtze River basin of China: patterns, threats and conservation. Biodivers. Conserv. 12, 1649–1685 (2003).

Guo, C., Li, W., Li, S. et al. Manipulation of fish community structure effectively restores submerged aquatic vegetation in a shallow subtropical lake. Environ. Pollut. 292, 118459 (2022a).

Guo, C., Li, S., Li, W. et al. Spatial variation in the composition and diversity of fishes inhabiting an artificial water supply lake, Eastern China. Front. Ecol. Evol. 10, 921082 (2022b).

IUCN. The IUCN Red List of Threatened Species. Version 2024-11 (2024).

Huang, L. & Li, J. Status of freshwater fish biodiversity in the Yangtze River Basin, China. Aquat. Biodivers. Conserv. Ecosyst. Serv. 13–30 (2016).

Jiang, H., Cheng, X., Geng, L., Tang, S., Tong, G. & Xu, W. Comparative study of the nutritional composition and toxic elements of farmed and wild *Chanodichthys mongolicus*. Chin. J. Oceanol. Limnol. 35, 737–744 (2017).

Jin, B. et al. Basin-scale approach needed for Yangtze River fisheries restoration. Fish Fish. 23, 1009–1015 (2022).

Kang, B., Huang, X., Yan, Y., Yan, Y. & Lin, H. Continental-scale analysis of taxonomic and functional fish diversity in the Yangtze River. Glob. Ecol. Conserv. 15, e00442 (2018).

Kitamura, J. Factors affecting seasonal mortality of rosy bitterling (*Rhodeus ocellatus kurumeus*) embryos on the gills of their host mussel. Popul. Ecol. 47, 41–51 (2005).

Knouft, J. H. Freshwater resources and COVID-19. In: Laituri, M., Richardson, R. B., Kim, J. (eds) The Geographies of COVID-19. Global Perspectives on Health Geography, Springer, Cham (2022).

Li, Q., Yao, M., Du, J., Chen, X., Li, M., Zhou, B. & Sun, R. Study on growth and development of early postembryonic stage of *Onychostoma sima*. Agric. Sci. Technol. 14, 354 (2013).

Li, W., Zhai, D., Wang, C., Gao, X., Liu, H. & Cao, W. Relationships among trophic niche width, morphological variation, and genetic diversity of *Hemiculter leucisculus* in China. Front. Ecol. Evol. 9, 691218 (2021).

Mei, Z. et al. The impact of fisheries management practices on the survival of the Yangtze finless porpoise in China. Aquat. Conserv. Mar. Freshw. Ecosyst. 29, 639–646 (2019).

Nabi, G., Hao, Y., Zeng, X. & Wang, D. Assessment of Yangtze finless porpoises (*Neophocaena asiaorientalis*) through biochemical and hematological parameters. Zool. Stud. 56 (2017).

Ren, K., Shen, X., Wang, M., Shi, X. & Liu, G. Wild training of captive *Spinibarbus denticulatus* juveniles. South China Fish. Sci. 16, 18–24 (2020).

Rahel, F. J. Homogenization of freshwater faunas. Annu. Rev. Ecol. Syst. 33, 291–315 (2002).

Wang, J.-P., Cheng, C.-Y., Ho, C.-W. & Ueng, Y.-T. Complete mitochondrial DNA genome of *Onychostoma sima* (Cypriniformes: Cyprinidae). Mitochondrial DNA 26, 135–136 (2015).

Xie, J. Y., Yan, Y., Yang, Y. H. & Lin, S. Q. Analysis on genetic structure of *Chanodichthys mongolicus* populations by mitochondrial COI gene sequences. Freshw. Fish. 49(3), 3–7 (2019).

Xu, G., Gu, R., Liu, H., Jiang, T., Du, F., Nie, Z., Yang, J. & Xu, P. Fluctuation of Sr/Ca in otoliths of *Coilia nasus* in the Yangtze River and the validation for the anadromous migratory history. J. Fish. China 38, 939–945 (2014).

Yang, T. & Percival, R. V. The emergence of global environmental law. Ecol. L. Q. 36, 615 (2009).

Yang, H. et al. Status of aquatic organisms resources and their environments in Yangtze river system (2017–2021). Aquac. Fish. (2023).

Yang, J. et al. A preliminary study on diet of the Yangtze finless porpoise using next-generation sequencing techniques. Mar. Mammal Sci. 35, 1579–1586 (2019).

Ye, S., Li, Z., Zhang, T., Liu, J. & Xie, S. Assessing fish distribution and threats to fish biodiversity in the Yangtze River Basin, China. Ichthyol. Res. 61, 183–188 (2014).

Yu, J., Xia, M., Guan, B., He, H., Chen, F. & Liu, Z. Interactive effects of bittering fish and mussel on the community structures of plankton and benthic macroinvertebrates. J. Lake Sci. 33, 1230–1240 (2021).

Yu, Z. T. Preliminary evaluation of the impact of large water conservancy projects on fish resources in the Yangtze River (in Chinese). J. Hydroecol. (2), 38–41 (1988).

Zhang, H., Kang, M., Shen, L. et al. Rapid change in Yangtze fisheries and its implications for global freshwater ecosystem management. Fish Fish. 21, 601–620 (2020).

Zhuang, H. & Wolf, S. A. Environmental public interest litigation: new roles for civil society organizations in environmental governance in China. Environ. Sociol. 7, 393–406 (2021).

